# Genome-wide approaches for the identification of markers and genes associated with sugarcane yellow leaf virus resistance

**DOI:** 10.1101/2020.09.04.283614

**Authors:** Ricardo José Gonzaga Pimenta, Alexandre Hild Aono, Roberto Carlos Villavicencio Burbano, Alisson Esdras Coutinho, Carla Cristina da Silva, Ivan Antônio dos Anjos, Dilermando Perecin, Marcos Guimarães de Andrade Landell, Marcos Cesar Gonçalves, Luciana Rossini Pinto, Anete Pereira de Souza

## Abstract

A major disease affecting sugarcane, a leading sugar and energy crop, is sugarcane yellow leaf (SCYL), caused by the sugarcane yellow leaf virus (SCYLV). Despite damages caused by SCYLV, the genetic basis of resistance to this virus remains largely unknown. Several methodologies have arisen to identify molecular markers associated with SCYLV resistance, which are crucial for marker-assisted selection and understanding response mechanisms to this virus. We investigated the genetic basis of SCYLV resistance using dominant and codominant markers and genotypes of interest for breeding. A sugarcane panel inoculated with SCYLV was analyzed for SCYL symptoms, and viral titer was estimated by RT-qPCR. This panel was genotyped with 662 dominant markers and 70,888 SNPs and indels with allele proportion information. We used polyploid-adapted genome-wide association analyses and machine-learning algorithms coupled with feature selection methods to establish marker-trait associations. While each approach identified unique marker sets associated with phenotypes, convergences were observed between them, demonstrating their complementarity. Lastly, we annotated these markers, identifying genes encoding emblematic participants in virus resistance mechanisms and previously unreported candidates involved in viral responses. Our approach could accelerate sugarcane breeding targeting SCYLV resistance and facilitate studies on biological processes leading to this trait.

## 1 Introduction

Sugarcane is one of the world’s most important crops, ranking first in production quantity and sixth in net production value in 2016^1^. It is by far the most relevant sugar crop, accounting for approximately 80% of the world’s sugar production^1–2^ and is also a prominent energy crop. However, it has an extremely complex genome; modern cultivars are the product of a few crosses between two autopolyploid species. *Saccharum spontaneum* (2*n* = 5x = 40 to 16x = 128; *x* = 8)^3^, a wild stress-resistant but low-sugar species, was hybridized and backcrossed with *Saccharum officinarum* (2*n* = 8x = 80, *x* = 10)^4^, which has a high sugar content but is sensitive to drought and susceptible to diseases. These procedures gave origin to plants with very large (ca. 10 Gb), highly polyploid, aneuploid and remarkably duplicated genomes^5–6^. This complexity directly affects sugarcane research and breeding; until recently, it prevented the use of codominance information in marker-assisted breeding strategies for this crop, limiting such approaches^7–8^.

One of the diseases that affect this crop is sugarcane yellow leaf (SCYL), which is caused by sugarcane yellow leaf virus (SCYLV), a positive-sense ssRNA virus belonging to the *Polerovirus* genus^9–10^. The expression of SCYL symptoms is complex and usually occurs in late stages of plant development, being mainly characterized by the intense yellowing of midribs in the abaxial surface of leaves^11–12^. SCYLV alters the metabolism and transport of sucrose and photosynthetic efficiency^13–14^, impairing plant development and eventually reflecting productivity losses^15–18^. Many SCYL symptoms may, however, be caused by other stresses or plant senescence^12,15,19^, making SCYL identification troublesome. Therefore, molecular diagnosis of SCYLV infection is of great importance; this was initially performed through immunological assays^11^, but more sensitive and sensible methods using reverse transcription followed by quantitative polymerase chain reaction (RT-qPCR) were later developed^20–22^.

Due to SCYL’s elusive symptomatology, SCYLV’s spread is silent; it is disseminated mostly during sugarcane vegetative propagation but is also transmitted by aphids, mainly the white sugarcane aphid *Melanaphis sacchari* (Zehntner, 1897)^11^. Unlike other pathogens, the virus is not efficiently eradicated by thermal treatments^23^; the only way to thoroughly eliminate it is by meristem micropropagation^24–25^, which is time-consuming and requires specialized infrastructure and personnel. These features make varietal resistance to SCYLV the most efficient resource to prevent damage and losses caused by this virus. Resistance has been explored in breeding programs and by a few genetic mapping studies^26–30^. However, research on SCYL genetics is not exempt from the difficulties generated by the complexity of the sugarcane genome^31^; due to this crop’s polyploid nature, most of these works employed dominantly scored molecular markers, implying a great loss of genetic information^32^. Additionally, they employed immunological methods to phenotype SCYLV resistance. The usage of dominant markers and the poor reliability of phenotyping were listed as key factors limiting the power of these studies^27–28^.

Here, we evaluated the efficacy of several genome-wide approaches to identify markers and genes associated with SCYLV resistance. We analyzed a panel of *Saccharum* accessions inoculated with SCYLV, which were graded for the severity of SCYL symptoms, and their viral titer was estimated by relative and absolute RT-qPCR. This panel was genotyped with amplified fragment length polymorphisms (AFLPs) and simple sequence repeats (SSRs), as well as single nucleotide polymorphisms (SNPs) and insertions and deletions (indels) obtained by genotyping-by-sequencing (GBS). We then employed three distinct methodologies to detect marker-trait associations: the fixed and random model circulating probability unification (FarmCPU) method using dominant AFLPs and SSRs; mixed linear modeling using SNPs and indels, in which allele proportions (APs) in each locus were employed to establish genotypic classes and estimate additive and dominant effects; and several machine learning (ML) methods coupled with feature selection (FS) techniques, using all markers to predict genotype attribution to phenotypic clusters. Finally, we annotated genes containing markers associated with phenotypes, discussing the putative participation of these genes in the mechanisms underlying resistance to SCYLV.

## 2 Results

### 2.1 Phenotypic Data Analyses

A total of 97 sugarcane accessions inoculated with SCYLV were evaluated for the severity of SCYL symptoms and for viral titer estimated by relative and absolute RT-qPCR quantification in two consecutive years, as comprehensively described in Supplementary Results. Based on best linear unbiased prediction (BLUP) estimations, symptom severity was not correlated with the viral titer determined by relative (p = 0.117) or absolute (p = 0.296) quantification. We found, however, a significant (p < 2.2e-16) and strong (r^2^ = 0.772) correlation between the values obtained by the two quantification methods, indicating their reliability (Supplementary Fig. 2).

Using BLUP values, we performed two hierarchical clustering on principal components (HCPC) analyses to investigate the classification of genotypes according to SCYLV resistance phenotypes – the first using BLUP values of SCYLV titers determined by RT-qPCR, and the second including BLUP values of all three traits analyzed. Both analyses indicated a division of the panel into three clusters (Supplementary Figs. 3-4) – named Q1-3 for the first HCPC and SQ1-3 for the second analysis. Factor maps wherein these groups are plotted onto the first two dimensions of HCPCs are shown in Fig. 1, and the attribution of genotypes to each cluster is available in Supplementary Table 4. Each group defined in the first HCPC presented significantly different SCYLV titers as estimated by both quantification methods (Supplementary Fig. 5, Supplementary Table 5). The second HCPC also resulted in a separation of groups with contrasting phenotypes: SQ1 accessions showed the least severe SCYL symptoms and the lowest titers of SCYLV; SQ2 accessions displayed significantly more severe disease symptoms and higher viral titers; and SQ3 accessions had the most severe disease symptoms and equally higher virus titers (Supplementary Fig. 6, Supplementary Table 5).

**Fig. 1.**
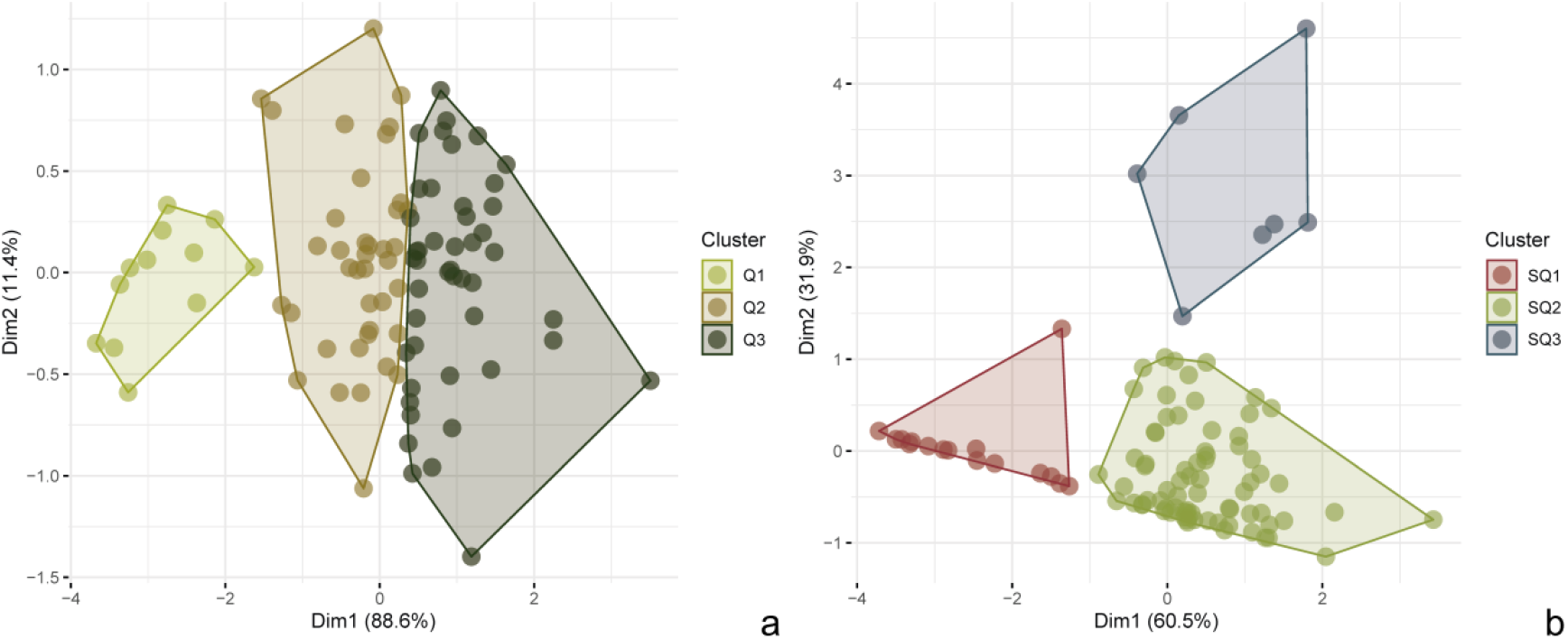
Factorial maps generated in the two hierarchical clustering on principal components (HPCP) analyses using BLUP values. (a) Factorial map of HCPCs performed using the SCYLV titer determined by RT-qPCR. A division into three clusters (Q1, Q2 and Q3) was considered. (b) Factorial map of HCPC performed using SCYL symptom severity and SCYLV titer determined by RT-qPCR. A division into three clusters (SQ1, SQ2 and SQ3) was considered

### 2.2 Genotyping and genetic analyses

After genotyping and filtering procedures, 93 accessions of the panel were successfully characterized with 550 AFLP fragments and 112 SSR fragments, totaling 662 polymorphic dominant markers. The GBS library constructed allowed the successful genotyping of 92 panel accessions, as described in detail in the Supplementary Results. We performed variant calling using BWA aligner and a monoploid chromosome set isolated from the *S. spontaneum* genome as a reference. This genome allowed the discovery of a large number of markers (38,710 SNPs and 32,178 indels) with AP information after rigorous filtering (Supplementary Tables 6-7). Additionally, unlike many of the references tested, it provided markers with information of position at chromosome level, allowing the estimation of long-distance linkage disequilibrium (LD). Pairwise LD between markers located within chromosomes was obtained and its decay was analyzed over distance. We observed high r^2^ values (∼0.4) between closely distanced markers, which dropped to 0.1 at approximately 2 Mb (Fig. 2).

**Fig. 2.**
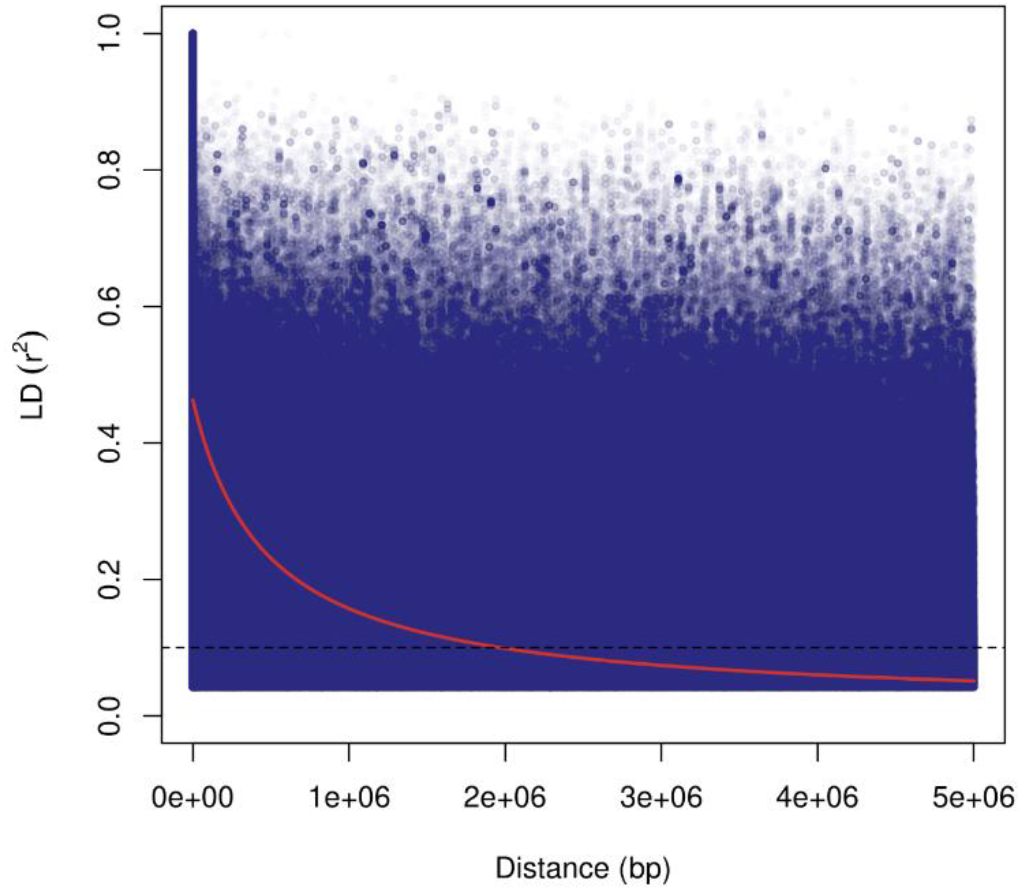
Decay of linkage disequilibrium (r^2^) as a function of physical distance (bp) between pairs of 67,007 single nucleotide polymorphisms (SNPs) and insertions and deletions (indels) located on *Saccharum spontaneum* chromosomes 1A-8A. Only r^2^ values with P < 0.05 are included

The genetic structure of the panel was investigated separately using the two marker datasets generated – AFLPs and SSRs scored as dominant and codominant SNPs and indels with AP information –, and three different approaches – a discriminant analysis of principal components (DAPC), a principal component analysis (PCA) followed by k-means and a Bayesian clustering implemented in STRUCTURE. Results are thoroughly described in the Supplementary Results, and Supplementary Table 8 summarizes the allocation of genotypes to the clusters identified in each analysis. Analyses performed with dominant markers identified two to four clusters, depending on the structure analysis employed (Supplementary Figs. 7-10); however, we observed extensive similarities between the groups identified in each method. A similar pattern was observed when the same three structure analyses were performed with codominant markers. Each method resulted in a unique separation of accessions, varying between two and three groups (Supplementary Figs. 11-14), but the clustering obtained by these different analyses was overall coincident. We found, however, that using dominant or codominant markers yielded noticeably different outcomes. Some overlap was observed between clusters identified by the analyses using each set of markers but, overall, groups identified by these analyses shared little resemblance. Additionally, the results from these methods did not present correspondences with the phenotype-based HCPCs.

### 2.3 Association Analyses

#### 2.3.1 FarmCPU

For FarmCPU analyses, we tested including matrices obtained from each genetic structure analysis as covariates and ran the models with no covariates. The distribution of the genomic inflation factor λ (Supplementary Fig. 15) was normal (p = 0.975) and no significant differences (p = 0.084) were observed between the inflation of p-values of models. Thus, we chose to conduct FarmCPU analyses using no covariates, as this resulted in the median value of λ closest to its theoretical value under the null hypothesis (λ = 1) and in appropriate profiles of inflation of p-values as seen in quantile-quantile (Q-Q) plots (Supplementary Fig. 16). Using a Bonferroni-corrected threshold of 0.05, one marker-trait association was detected for symptom severity and five associations were detected for the viral titer estimated by each quantification method – with one marker being mutually associated with both. The percentage of phenotypic variance explained by each marker ranged from 9 to 30% (Supplementary Table 9).

#### 2.3.2 Mixed Modeling

Twelve combinations of population structure (Q) and kinship (K) matrices were tested as effects in the codominant association models. The distribution of λ in each Q + K combination (Supplementary Fig. 17) was not normal (p = 3.253e-06) and no significant differences (p = 0.869) were detected between models. Thus, following analyses were conducted with a Q + K combination that resulted in the median value of λ closest to 1, which was obtained with the combination of the first three PCs from a PCA with both the realized relationship (MM^T^) and pseudodiploid kinship matrices. As the MM^T^ matrix is directly computed by the GWASpoly package, we considered the Q_PCA_ + K_MM_ combination to be the most straightforward. Q-Q plots of the association analyses for SCYL symptom severity and SCYLV relative and absolute quantifications can be found in Supplementary Fig. 18; in general, all models showed appropriate inflation of p-values.

A stringent significance threshold (p < 0.05 corrected by the Bonferroni method) was used to identify 35 nonredundant markers significantly associated with SCYL symptom severity (Fig. 3). Using this correction, no markers were significantly associated with SCYLV titer. In an attempt to establish a less conservative threshold for association analyses of these two traits, we employed the false discovery rate (FDR) for the correction of p-values, which resulted in very low significance thresholds and the identification of thousands of associations as significant. Therefore, we ultimately opted to use an arbitrary threshold of p < 0.0001 to determine markers strongly associated with the two quantification traits. This resulted in 13 and 9 markers associated with SCYLV titer determined by relative and absolute quantifications, respectively (Fig. 3); one marker was common to both analyses. Supplementary Table 10 supplies information on all marker-trait associations identified by this approach. For each trait, we observed a redundancy between markers identified as significant by different marker-effect models; this observation was particularly common between the simplex dominant alternative and the diploidized models.

**Fig. 3.**
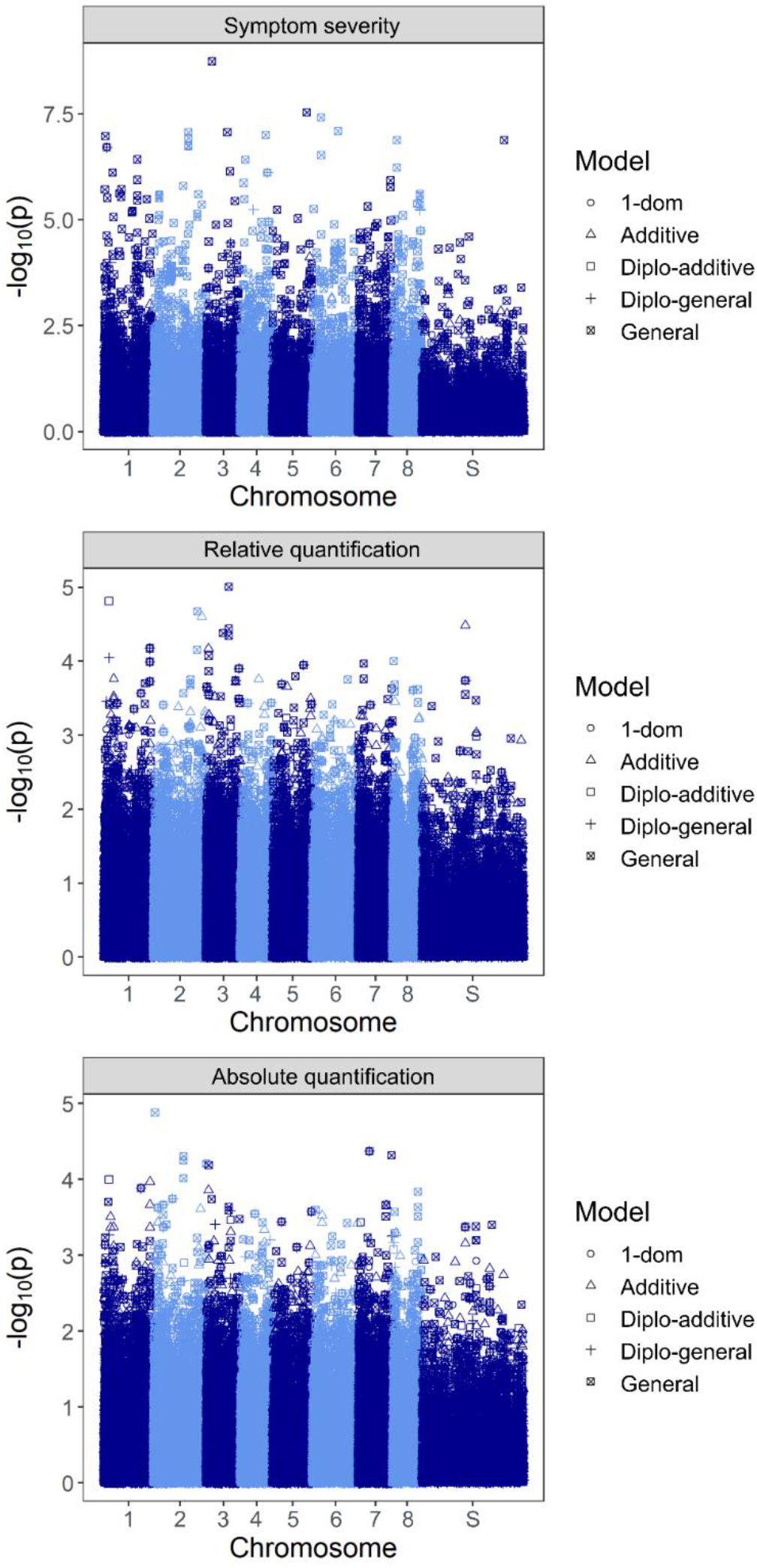
Manhattan plots generated in association analyses using the best linear unbiased predictor (BLUP) values of the three traits analyzed. Six different models were tested: general, additive, simplex dominant reference (1-dom-ref), simplex dominant alternative (1-dom-alt), diploidized general (diplo-general) and diploidized additive (diplo-additive). On the x-axis, S represents scaffolds not associated with any of the *S. spontaneum* chromosomes

#### 2.3.3 Machine Learning Coupled with Feature Selection

As a last marker-trait association method, we tested eight ML algorithms for predicting the attribution of genotypes to the phenotypic clusters identified in the HCPCs. When assessing their potential in this task using the full marker dataset, predictive accuracies varied greatly depending on the method and phenotypic groups under analysis. They were lower for the prediction of clusters associated with viral titer (Q), ranging between 39.2-49.6%, with an average of 44.5% (Supplementary Fig. 19a). For clusters identified including symptom severity data (SQ), accuracies were overall higher, albeit varying even more and being still unsatisfactory; they ranged between 7.9-73.9% (Supplementary Fig. 19b) and had an average of 58%. Therefore, we tested applying five FS methods to reduce the marker dataset, and constructed three additional reduced marker datasets consisting of intersections between FS methods.

These procedures led to considerably higher accuracies in predicting Q and SQ clusters. Three FS methods (FS1, FS2 and FS4) presented notably superior effects in increasing accuracy in both cases (Supplementary Fig. 20). In the two scenarios, the most accurate model-FS combination was a multilayer perceptron neural network (MLP) coupled with FS2, which was composed of 232 markers for Q and 170 markers for SQ. This combination resulted in average accuracies of 97.6% and 96.5% for the prediction of Q and SQ, respectively (Supplementary Tables 11 and 12). However, in both scenarios, MLP achieved the second-best results when using Inter2 datasets, composed by markers present in at least two out of the three best FS methods; these represented 190 markers for Q and 120 markers for SQ. With this strategy, we could achieve equally high accuracies (95.7% for Q and 95.4% for SQ) with further reductions in marker numbers. To farther evaluate the performance of MLP, we produced receiver operating characteristic (ROC) curves and calculated their respective area under the curves (AUCs). Prior to FS, MLP did not present satisfactory results, with ROC curves very close to the chance level and AUCs of 0.45-0.61 for Q and 0.40-0.56 for SQ (Fig. 4a). When Inter2 was used, ROC curves showed much better model performances, with AUCs of 1.00 for Q and of 0.98-1.00 for SQ (Fig. 4b). These results confirm that Inter2 markers are in fact associated with SCYLV resistance and that MLP is an appropriate model to predict clustering based on this dataset. The markers representing the reduced datasets associated with Q and SQ clusters can be found in Supplementary Tables 13 and 14, respectively. We observed twelve marker overlaps between the two datasets; interestingly, several of these markers were also identified as associated with phenotypes in the FarmCPU and mixed modeling analyses.

**Fig. 4.**
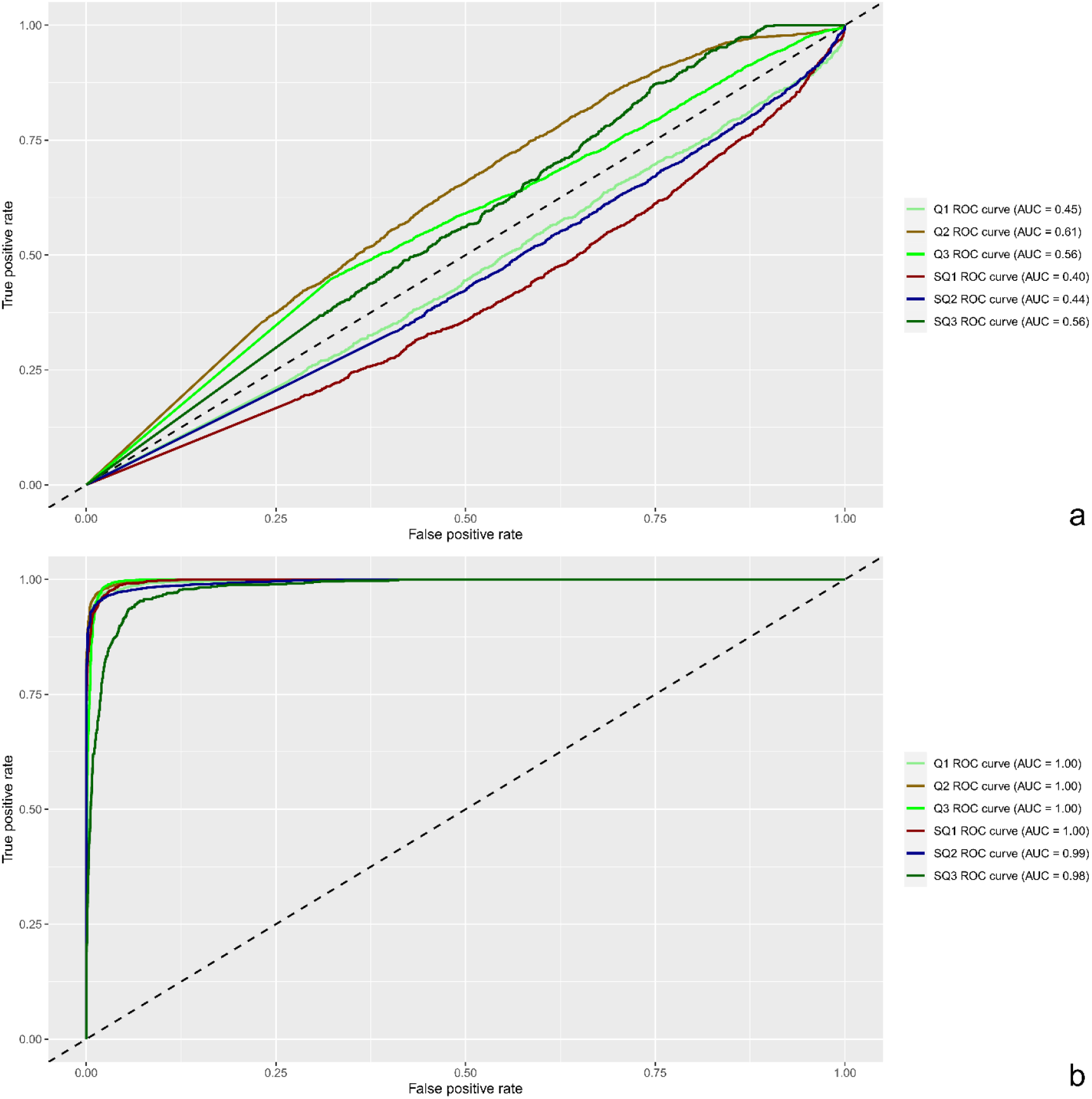
Receiver operating characteristic (ROC) curves and area under the curve (AUC) results regarding the performance of MLP in predicting clustering by by SCYLV titer determined by RT-qPCR (Q) and SCYLV titer determined by RT-qPCR and SCYL symptom severity (SQ). (a) Model performance obtained using the full marker dataset. **(b)** Model performance obtained using the marker dataset obtained from the intersection of at least three of the three best feature selection methods employed in the study (Inter2)

### 2.4 Marker Mapping and Annotation

For a better visualization of the physical location of all markers associated with SCYLV resistance, we constructed a map of their distribution along *S. spontaneum*’s “A” chromosomes (Fig. 5), in which we also included markers identified as associated with SCYLV resistance in previous mapping studies. Overall, markers were considerably spread along chromosomes; however, we observed regions of dense concentration of markers identified by various methods, such as the long arms of chromosomes 1 and 3. We also verified the proximity between several markers identified in the present work and by other authors, indicating their convergence and the reliability of the methods employed here.

**Fig. 5.**
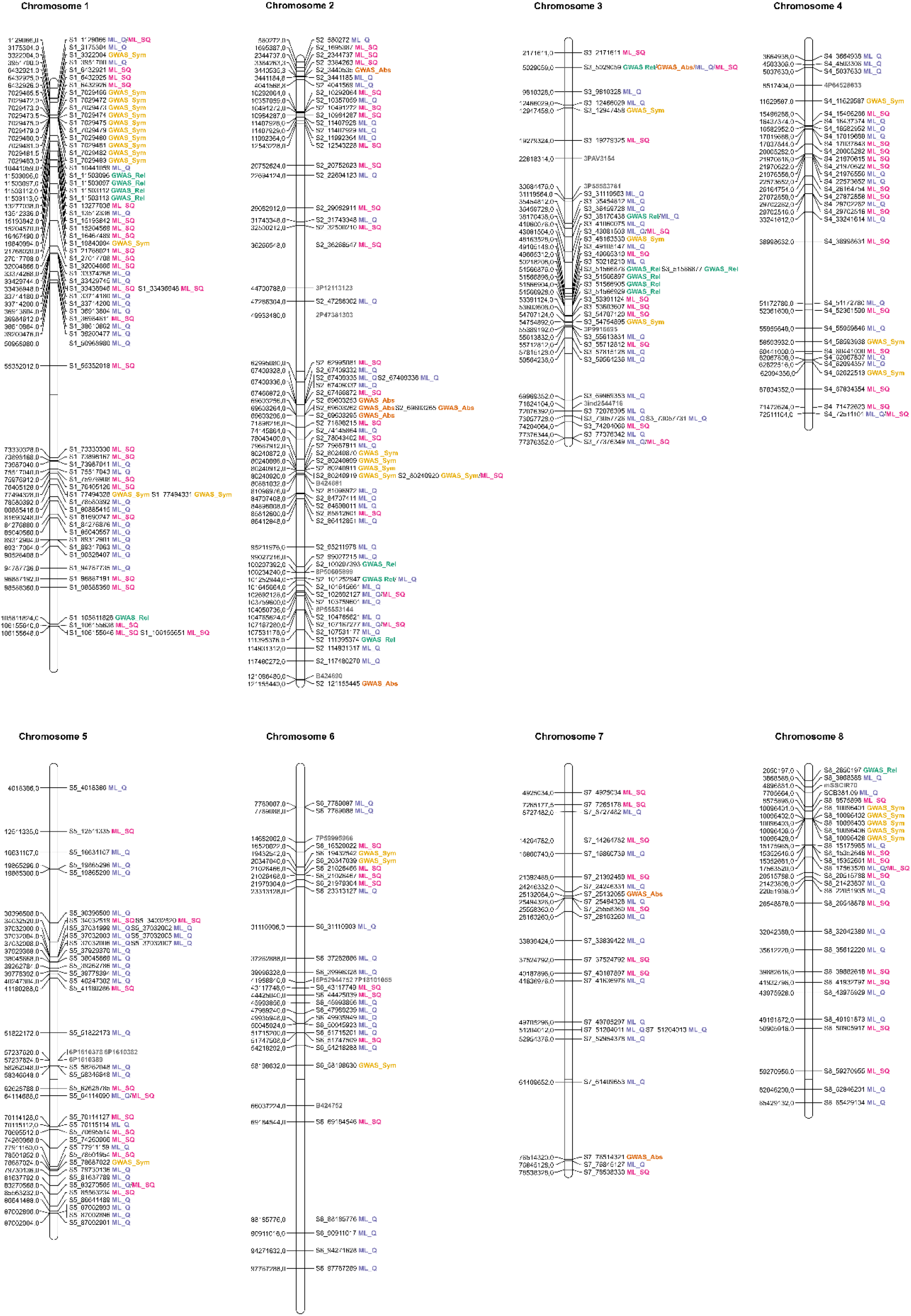
Distribution of markers associated with SCYLV resistance along *Saccharum spontaneum* “A” chromosomes. In each chromosome, marker positions are shown on the left, and marker names are indicated on the right, labeled and colored according to the method employed for their identification. Markers identified by previous mapping studies are colored in gray

Out of the 362 nonredundant markers associated with all phenotypes, 176 were located in genic regions and could be annotated by aligning their 2000-bp neighboring regions with the coding sequences (CDSs) of 14 Poaceae species and *A. thaliana* genomes; Supplementary Table 15 contains data on the alignment with the highest percentage of identity for each marker. In some cases, where two or more markers were closely located, coincident alignments and annotations were obtained; consequently, 148 genes were representative of all the best alignments. The large majority of top-scoring alignments (117) occurred with CDSs of *Sorghum bicolor*, the phylogenetically closest species among those used for alignment; fewer alignments also occurred with the CDSs of other species. Several of the annotated genes could be associated with plant resistance to viruses, as detailed in the discussion.

## 3 Discussion

We evaluated the severity of SCYL symptoms and SCYLV titer in a panel of 97 sugarcane accessions. These two traits are of great concern to breeding, as both have been associated with higher yield losses in SCYLV-infected sugarcane plants^18,22,33^. Prior to phenotyping, plants were subjected to high and uniform SCYLV inoculum pressure, an innovation over all previous SCYLC genetic mapping studies^26–30^, which relied on natural infection under field conditions. Using RT-qPCR, currently regarded as the most precise method for SCYLV quantification^22^, we assessed the viral titer in these genotypes. We found a strong and positive correlation between the BLUPs calculated for the SCYLV titers obtained by the two quantification methods employed, showing the consistency of the data. The absence of a perfect correlation might have arisen from intrinsic differences between methods, which have been responsible for disparities in viral quantification by RT-qPCR in other plant-virus interactions^34^.

However, we observed no quantitative correlation between the severity of SCYL symptoms and SCYLV titers across the sugarcane genotypes analyzed. This finding corroborates a growing body of evidence suggesting that these traits are not strongly or necessarily correlated, i.e., high SCYLV titers are not a guarantee of more severe yellowing or of its development at all^35–37^. This reinforces the importance of SCYLV molecular screening of sugarcane clones by breeding programs; this should be done to avoid the employment of genotypes that accumulate high viral loads asymptomatically but may inconspicuously suffer yield losses, in addition to serving as a virus reservoir for vector transmission to other susceptible genotypes.

To further explore this issue, we performed two HCPC analyses to discriminate accessions based on their response to SCYLV, which led to the separation of clusters with considerable phenotypic differences. In the first HCPC, using only viral quantification data, we could discern groups with significant variation in viral titers; in the second analysis, which also included symptom severity data, clusters with even more contrasting responses to SCYLV could be discriminated. In 1983, Cooper and Jones^38^ proposed a terminology addressing plant responses to viral infections that is still employed today^39–41^. According to this proposal, once infected, plants present differences in the ability to restrict viral replication and invasion; at the extremes of a spectrum of behaviors are plants termed susceptible and resistant. Additionally, they may also respond differently to the infection in terms of symptom development: another spectrum exists, at the extremes of which are sensitive and tolerant plants. In view of this nomenclature, we propose that the clusters identified in this second HCPC be described as follows: (SQ1) resistant, for sugarcane genotypes distinguished by low SCYLV titer and mild or no SCYL symptoms; (SQ2) tolerant, for genotypes that, despite exhibiting higher viral titers, presented few or no disease symptoms; and (SQ3) susceptible, for genotypes with the most severe symptoms and presenting high viral titers. This classification per se is of great use in sugarcane breeding, as it distinguishes not only sources of tolerance to SCYLV but also an exceptionally promising group of truly resistant genotypes.

Our main objective was, however, to identify markers associated with SCYLV resistance in a broader sense. With this aim, we performed genotyping with a combination of dominant and codominant markers, which has never been described for sugarcane. We evaluated the impact of using genomic references from various backgrounds in variant calling from GBS. In previous sugarcane GWASs, this was performed using the genome of *S. bicolor*^30,42–44^, a close relative species with a well-assembled and annotated genome; however, in our analyses, this reference yielded a number of markers considerably inferior to other references. The methyl-filtered genome of the SP70-1143 cultivar yielded the most markers, in agreement with a previous study employing GBS^45^; this is a plausible outcome, as this method avoids sampling of methylated regions^46^ which were also filtered out for this genomic assembly^47^. However, to choose the best reference for further analyses, we also considered the quality of the assembly, which greatly affects the results of GWASs in polyploids^48^. The best-assembled sugarcane genome available to date is the allele-defined genome of a haploid *S. spontaneum* accession^49^. Despite presenting one of the highest total tag alignment rates, this reference also gave a very high rate of multiple alignments, leading to the identification of relatively few markers. This was probably due to the alignment of tags to homeologous regions of different alleles rather than to the duplicated regions that we intended to avoid. To circumvent this situation, we conducted our analyses with markers isolated using a monoploid chromosome set obtained from this genome, which provided a large number of markers with reliable position information.

Using these codominant markers, we analyzed the decay of LD over distance. LD has long been hypothesized to be high in sugarcane due to the short breeding history and narrow genetic basis of this crop; many studies using dominant markers have estimated it to be especially high at 5-10 cM^50–54^. The first study to use SNPs for this task and estimate LD decay in bp^55^ indicated that LD was extremely long lasting, with the average r^2^ decaying to 0.2 at 3.5 Mb in hybrids. Our results further confirm the persistence of LD at long distances in sugarcane, albeit indicating that it decayed more quickly – with r^2^ dropping to 0.2 at less than 1 Mb and to 0.1 at 2 Mb. These results impact mapping studies, as a high LD implies that a low density of markers might be needed for accurate mapping of quantitative traits.

We tested several approaches to evaluate population structure in the panel using each distinct marker dataset generated, which yielded remarkably different results. Studies contrasting the usage of dominant and codominant markers in plants have shown discrepancies in measures of genetic structure and diversity^56–58^, but this sort of comparison has never been performed including markers with dosage information in polyploids – let alone in sugarcane. In this crop, the most relatable findings available are those reported by Creste et al.^59^, who showed that using different dominant markers can bias genetic analyses, and thus the choice of marker must be guided by the specific goal of each study. For GWASs – for which a high density of markers is usually necessary – SNPs and indels are currently more cost-effective, as they can be easily identified in much larger numbers, in addition to offering the possibility of estimating highly-informative allele dosages or APs^60–62^. Hence, we believe the results we obtained with codominant SNPs and indels are more reliable, as they lean on much more genetic information.

In contrast with the differences arising from the type of marker used, we observed little divergence between results of different structure methods performed with each marker dataset, and eventual discrepancies did not result in significant differences in the inflation of the association models, whose patterns were similar to those of previous studies^30,43–44,54^. Therefore, we opted to perform association analyses using the covariates that resulted in the value of λ closest to 1. For FarmCPU, this corresponded to the “naive” model with no covariates; for codominant mixed modeling analysis, this was the Q_PCA_ + K_MM_ combination. K_MM_ is the usual choice of relationship matrix in polyploid association mapping^63–65^, as Q matrices obtained from PCA are commonly used to control population structure in GWASs^66–68^.

FarmCPU analyses using dominant markers identified one AFLP fragment significantly associated with symptom severity, which explained a small part of the phenotypic variation (r^2^ = 0.116). Eight out of the nine markers associated with viral titer explained larger parts of the variation in the phenotypes (21-30%). These results are more promising than those obtained in a previous dominant GWAS targeting SCYLV resistance, which found r^2^ ranging between 0.09-0.14^27^. Albeit low, values in this range are very common in sugarcane association studies. Evidence indicates that almost all of this crop’s traits are highly quantitative, with the notable exception of brown rust resistance^69–70^. For other relevant traits, it is common to find the most associated markers explaining ≤ 10% of the phenotypic variation^28,42,54^.

A few authors have suggested that these suboptimal results could be improved with the usage of markers with dosage; this was also performed here, using SNPs and indels with AP information. Although codominant mixed modeling analyses successfully identified markers associated with SCYL symptom severity using the Bonferroni correction, the same was not observed for SCYLV titer. This was probably influenced by the modest size of the panel, a factor that restricts the power of GWASs^71–72^. As previously noted by Racedo et al.^73^, assembling and phenotyping large sugarcane association panels is a challenging task; thus, it is not uncommon for association studies of this crop to evaluate fewer than 100 genotypes^42,73–76^. Our study was particularly burdensome, as extremely laborious inoculation and quantification techniques were employed to generate highly reliable phenotypic data. Furthermore, the Bonferroni method is notorious for its conservative nature, poorly controlling false negatives^77–79^. This led us to establish an arbitrary threshold (p < 0.0001) to select markers strongly associated with SCYLV titer for further investigation. Using this methodology, we identified 57 nonredundant markers associated with the three phenotypes.

As a last approach to identify marker-trait associations, we tested several ML algorithms coupled with FS methods to predict genotype attribution to phenotypic clusters identified by HCPC analyses. Unlike methods built on classical statistics, these algorithms are not as heavily impacted by the sample size. We could achieve very high accuracies of prediction (up to 95%) with considerably reduced datasets comprising 120-190 markers. These results are very similar to what was obtained for predicting sugarcane brown rust resistance groups, where an accuracy of 95% was obtained using 131 SNPs^62^. Marker datasets selected by ML have rarely been employed in genetic association studies in plants, but the few existing examples show their power to identify genes associated with phenotypes of interest^80–82^.

We annotated 176 markers associated with SCYLV resistance to 148 genes. Many candidates do not allow extensive discussion on their involvement in resistance to this disease, as they either have very generic descriptions or have not been previously linked to plant virus resistance. Other proteins have occasionally been associated with responses to viruses but are members of very large gene families with extremely diverse biological roles and will not be discussed. Remarkably, a few candidates encode proteins previously associated with the response to SCYLV infection; this was the case for SbRio.10G317500.1, encoding a peroxidase precursor. Peroxidases are long known to be activated in response to pathogens, but most notably, a guaiacol peroxidase has been shown to be more active in sugarcane plants exhibiting SCYL symptoms than in uninfected or asymptomatic plants^83^. Our results provide further evidence that these enzymes are in fact involved in the response to SCYLV. Other candidates harboring markers associated with SCYLV resistance encode proteins with motifs previously associated with SCYLV resistance^30^: Sobic.001G023900, encoding a GATA zinc finger protein, and Sobic.001G200200 and Zm00001d037864_T030, which encode proteins containing tetratricopeptide repeats.

Other annotations included classic participants in more general disease resistance mechanisms, such as several genes encoding proteins with leucine-rich repeat (LRR) motifs. These structures are part of nucleotide-binding LRR (NBS-LRR) proteins, receptors that detect pathogen-associated proteins and elicit effector-triggered immunity^84^, having been shown to determine resistance to viruses in plants^85–87^. We found two LRR proteins (Sobic.008G156600.1 and Sobic.001G452600.1), one disease resistance NBS-LRR (Sobic.007G085400.1) and one N-terminal leucine zipper NBS-LRR resistance gene analog (Sobic.005G203500.1) associated with SCYLV resistance. Furthermore, we annotated one gene (Sobic.009G204800.1) that encodes a precursor of a receptor-like serine/threonine-protein kinase, i.e., the family to which LRR proteins belong. Yang et al.^30^ also identified a serine/threonine-protein kinase associated with SCYLV resistance. We consider these proteins highly promising candidates to be involved in the recognition of infection by SCYLV, which could trigger response mechanisms leading to the restriction of the virus. Further virus-host interaction studies involving these proteins might help confirm this hypothesis, which would represent a major breakthrough in understanding resistance to SCYLV.

Two other annotated genes were readily identified as involved in plant disease resistance mechanisms. Sobic.010G131300.2 contains a Bric-a-Brac, Tramtrack, Broad Complex/Pox virus and Zinc finger (BTB/POZ) domain, while Sobic.007G198400.1 contains two BTB domains, as well as ankyrin repeat regions. These domains are present in and are essential for the function of NONEXPRESSOR OF PATHOGENESIS-RELATED GENES 1 (NPR1), a central player in plant disease responses^88–89^. This family of transcription factors is involved in establishing both systemic acquired resistance and induced systemic resistance^90^, mediating the crosstalk between salicylic acid and jasmonic acid/ethylene responses^91^. Correspondingly, NPR1 has been widely shown to be involved in resistance to viruses^92–93^; therefore, it is reasonable to suggest its participation in the response to infection by SCYLV.

We also found a few candidates with putative roles in the RNA interference mechanism, one of the most prominent processes that contribute to resistance against viruses in plants. This is the case for Sobic.001G214000.1, which encodes a Dicer. Dicers are part of a mechanism known as RNA silencing, recognizing and cleaving long double-stranded RNA molecules into mature small RNAs that guide the cleavage of viral mRNAs and disrupt virus replication^94^; accordingly, these nucleases have been linked to resistance to viruses in several plant species^95–96^.

Another gene possibly involved in RNA interference is Sobic.009G121100, encoding a protein related to calmodulin binding – a calcium transducer that regulates the activity of various proteins with diverse functions^97^ and has been widely implicated in viral resistance in plants, often playing roles in RNA interference^98–100^. Therefore, we consider these genes promising candidates in the regulation of SCYLV replication and spreading *in planta*, as well as in the development of SCYL symptoms.

Two additional annotations linked to the mechanism of RNA interference are those of genes encoding proteins with F-box domains, SbRio.03G158900 and Sobic.002G019750.1. F-box proteins are involved in virus resistance in several plant species^101–102^; a particularly interesting case is FBW2 from *A. thaliana*, which regulates AGO1, an Argonaute protein with a central role in RNA silencing^103^ and repression of target viral RNAs^104–106^. Even more intriguing is the fact that one of the proteins encoded by the SCYLV genome, P0, contains an F-box-like domain and mediates the destabilization of AGO1, leading to the suppression of host gene silencing^107^. Whether the F-box proteins identified here play active roles in silencing of SCYLV remains a question to be investigated by further studies.

Other annotated genes may represent host factors involved in various steps of plant-virus interactions. For instance, Sobic.010G160500.4 encodes an RNA helicase with a DEAD-box domain, which are often coopted by viruses to promote viral translation or replication, playing important roles in regulating infection^108–110^. Similarly, soluble N-ethylmaleimide-sensitive-factor attachment protein receptor (SNARE) proteins such as Sobic.001G528000.1 are essential in the biogenesis and fusion of vesicles of several plant viruses^111–114^. We also found one gene encoding a myosin (Sobic.002G108000.1) and two genes related to kinesin (Sobic.001G346600.1 and Sobic.001G399200.2), filament-associated motor proteins involved in the transport of organelles^115^. In a few cases, both myosins^116–118^ and kinesins^119^ have been shown to be involved in viral intercellular movement through poorly understood mechanisms. One last interesting annotation was Sobic.003G101500.1, a protein with a DNAJ domain. DNAJs have been shown to interact with proteins of various plant viruses and to be associated with resistance, sometimes being crucial for virus infection and spread^120–123^. We consider these genes to be promising candidates as host cofactors in the response to SCYLV infection.

In conclusion, this array of genome-wide analyses allowed us to detect markers significantly associated with SCYLV resistance in sugarcane. If validated, these markers represent an especially valuable resource for sugarcane breeding programs, as the results can be directly employed in marker-assisted strategies for the early selection of clones. The annotation of several genes wherein these markers are located revealed many candidates with long-established and pivotal roles in viral disease resistance, further demonstrating the efficiency of the methods employed for this purpose. Additionally, this annotation provides valuable insights into the unexplored mechanisms possibly involved in sugarcane’s response to infection by SCYLV, introducing new candidates whose role in this process can be further investigated in future studies.

## 4 Material and Methods

### 4.1 Plant Material and Inoculation

The plant material and inoculation methods employed in the present study are described by Burbano et al.^124^. The experimental population consisted of a panel of 97 sugarcane genotypes comprising wild germplasm accessions of *S. officinarum*, *S. spontaneum* and *Saccharum robustum*; traditional sugarcane and energy cane clones; and commercial cultivars originating from Brazilian breeding programs (Supplementary Table 1). To ensure plant infection with SCYLV, a field nursery was established in March 2016 at the Advanced Centre for Technological Research in Sugarcane Agribusiness located in Ribeirão Preto, São Paulo, Brazil (4°52’34” W, 21°12’50” S). Seedlings from sprouted setts of each genotype were planted in 1-meter plots with an interplot spacing of 1.5 meters. The cultivar SP71-6163, which is highly susceptible to SCYLV^15^, was interspersed with the panel genotypes. *M. sacchari* vector aphids were reared on RT-PCR tested SCYLV-infected SP71-6163 plants; after an acquisition access period of at least 48 hours, aphids were released weekly in the field nursery in July 2016. After plant growth, setts obtained from this nursery were used to install a field experiment following a randomized complete block design with three blocks in May 2017. Plants were grown in 1-meter-long three-row plots with row-to-row and interplot spacings of 1.5 and 2 meters, respectively. Each row contained two plants, totaling six plants of each genotype per plot. To further assist infection by SCYLV, the cultivar SP71-6163 was planted in the borders and between blocks, and *M. sacchari* aphids were again released in the field weekly for five months, starting from November 2017.

### 4.2 Phenotyping

Plants were phenotyped in two crop seasons: plant cane in June 2018 and ratoon cane in July 2019. The severity of SCYL symptoms was assessed by three independent evaluators, who classified the top visible dewlap leaves (TVDLs) of each plot using a diagrammatic scale established by Burbano et al.^124^, as shown in Supplementary Fig. 1. In the same week as symptom evaluation was performed, fragments from the median region of at least one TVDL per plot were collected and stored at −80°C until processing. Total RNA was extracted from this tissue using TRIzol (Invitrogen, Carlsbad, USA). Samples were subjected to an additional purification process consisting of three steps: (i) mixing equal volumes of RNA extract and chloroform, (ii) precipitating the RNA overnight with 2.5 volumes of 100% ethanol and (iii) a conventional cleaning step with 70% ethanol. RNA was then quantified on a NanoDrop 2000 spectrophotometer (Thermo Scientific, Waltham, USA) and subjected to electrophoresis on a 1% agarose gel stained with ethidium bromide for integrity checks. Samples were next diluted, treated with RNase-Free RQ1 DNase (Promega, Madison, USA), quantified and diluted again for standardization, and converted to cDNA using the ImProm-II Reverse Transcription System kit (Promega, Madison, USA).

The SCYLV titer in each sample was determined by qPCR using GoTaq qPCR Master Mix (Promega, Madison, USA) on a Bio-Rad CFX384 Touch detection system (Bio-Rad, Philadelphia, USA). Two viral quantification methodologies were employed – one relative and one absolute – using primers and conditions as described by Chinnaraja and Viswanathan^125^. For both methods, a set of primers was used to amplify a 181-bp fragment from SCYLV ORF3 (YLSRT). For the relative quantification, an additional set of primers was used to amplify a 156-bp fragment of the 25S subunit of sugarcane ribosomal RNA (25SrRNA), used as an internal control. The 2^−ΔΔCT^ method^126^ was used to correct cycle threshold (CT) values; the sample with the highest CT and a melting temperature of 82.5 ± 0.5°C for the YLSRT primers was used as a control for phenotyping in each year. The absolute quantification followed the methodology described by Chinnaraja et al.^37^. A pGEM-T Easy vector (Promega, Madison, USA) cloned with a 450-bp fragment from SCYLV ORF3 previously amplified by RT-PCR was used to construct a serial dilution curve with six points and tenfold dilutions between points, which was amplified on all qPCR plates. All reactions were performed using three technical replicates.

### 4.3 Phenotypic Data Analyses

The normality of phenotypic data was assessed by Shapiro-Wilk tests, and normalization was carried out using the bestNormalize package^127^ in R software^128^. BLUPs were estimated for each trait with the breedR R package^129^ using a mixed model as follows:

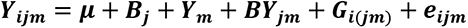

where *Yijm* is the phenotype of the i^th^ genotype considering the j^th^ block and the m^th^ year of phenotyping. The trait mean is represented by *μ*; fixed effects were modeled to estimate the contributions of the j^th^ block (*B_j_*), the m^th^ year (*Y_m_*) and the interaction between block and year (*B_Yjm_*). Random effects included the genotype (*G*) and the residual error (*e*), representing nongenetic effects.

Pearson’s correlation tests were performed using the BLUPs to check the correlation between traits, and correlation distributions were plotted using the GGally R package^130^. To investigate the separation of genotypes according to phenotypes, we performed two HCPC analyses with the factoMineR package^131^ – first using only viral quantification and then employing the three analyzed traits. The factoextra R package^132^ was used to plot graphs associated with these analyses. Statistical differences between the phenotypes of the clusters identified in each HCPC were assessed by Kruskal-Wallis tests or analyses of variance (ANOVAs), depending on the distribution of the data. Post hoc Dunn’s tests using the Bonferroni correction were performed with the R package dunn.test^133^ to verify pairwise differences between clusters.

### 4.4 Genotyping

#### 4.4.1 Dominant Markers

Total DNA was extracted from leaves of each genotype following the method described by Aljanabi et al.^134^. AFLPs were developed using *Eco*RI and *Msp*I restriction enzymes (New England BioLabs). Digestion reactions were prepared in a final volume of 20 μL containing 300 ng DNA, 2.5 U of each restriction enzyme in 1X RL Buffer (New England BioLabs) and incubated for 3 hours at 37°C and for 5 min at 70°C. Adapter ligation was conducted in a final volume of 40 μL containing 20 μL of the digestion reaction, 5 X buffer (40 mM Tris pH 8.4, 100 mM KCl), 0.5 μM *Eco*RI adaptor, 5 μM *Msp*I adaptor, 1 mM ATP and 0.85 U of T4 DNA ligase (67 U/μL) (New England BioLabs). Ligation was performed at 37°C for 2 hours and 16°C for 16 hours. Preamplification was conducted with primers complementary to restriction enzyme adaptors and devoid of selective nucleotides at the 3’ end (*Eco*RI+0 and *Msp*I+0 primers) and using a 6X dilution of the digestion/ligation product. This reaction was performed in a final volume of 15 μL containing 2 μL of the 6X dilution digestion/ligation product, 1X PCR buffer (20 mM Tris pH 8.4, 50 mM KCl), 3.3 μM *Eco*RI+0 and *Msp*I+0 primers, 0.17 mM dNTPs, 2 mM MgCl_2_ and 0.07 U Taq DNA polymerase. The cycling conditions were as follows: 29 cycles at 94°C for 30 seconds, 56°C for 1 minute and 72°C for 1 minute. Preamplification reactions were diluted 10X and used for selective amplification reactions using combinations of *Eco*RI/*Msp*I primers with three selective nucleotides at the 3’end and the *Eco*RI primer labeled with fluorophores IRDye700 or IRDye800. Thirty-five selective primer combinations were used (Supplementary Table 2). The reaction was performed in a final volume of 10 μL containing 2.5 μL of the 10X diluted preamplification, 1X PCR buffer (20 mM Tris pH 8.4, 50 mM KCl), 0.05 μM of selective Eco700 labeled primer (or 0.07 μM Eco800 primer), 0.25 μM for Msp selective primer, 0.25 μM dNTPs, 2 mM MgCl_2_, 0.5 U of Taq DNA polymerase. Cycling conditions were as follows: 94°C for 30 seconds, 65°C for 30 seconds and 72°C for 1 minute followed by 12 cycles at 94°C for 30 seconds, 65°C for 30 seconds (decreasing 0.7°C/cycle) and 72°C for 1 minute, followed by 23 cycles of 94°C for 30 seconds, 56°C for 30 seconds and 72°C for 1 minute. Final amplicons were separated on a 6% denaturing polyacrylamide gel and visualized with a LI-COR 4300 DNA Analyzer (LI-COR, Lincoln, NE, USA).

Twelve SSR loci previously isolated from the sugarcane expressed sequence tag database^135–138^ were used for SSR genotyping (Supplementary Table 3). PCR mixes were prepared and amplifications were conducted in a Bio-Rad MyCycler thermocycler (Bio-Rad, Philadelphia, USA) following the conditions previously established by Oliveira et al.^137^ and Marconi et al.^138^; primers were labeled with fluorescent dyes IRDye700 and IRDye800 to allow band visualization. Amplicons were separated on a 6% denaturing polyacrylamide gel and visualized with a LI-COR 4300 DNA Analyzer. Due to sugarcane polyploidy, both AFLPs and SSRs were treated as dominant and scored based on the presence (1) or absence (0) of bands. After genotyping, genotypes and markers with over 10% missing data were removed, as well as markers with a MAF below 10%.

#### 4.4.2 Genotyping-by-sequencing

Genomic DNA was extracted from leaves using the GenElute Plant Genomic DNA Miniprep Kit (Sigma-Aldrich, St. Louis, USA). The integrity of the DNA was verified by electrophoresis on a 1% agarose gel stained with ethidium bromide, and its concentration was determined using a Qubit 3.0 fluorometer (Thermo Scientific, Wilmington, USA). The construction of the GBS library was based on a protocol by Poland et al.^139^ and used a combination of *Pst*I and *Mse*I restriction enzymes. For operational reasons, 94 out of the 97 genotypes of the panel were included in the library, which did not include genotypes 87, 88 and 95 (see Supplementary Table 1). The library was subjected to a purification step using polyethylene glycol as described by Lundin et al.^140^ with slight modifications. It was then validated with a Fragment Analyzer (Agilent Technologies, Santa Clara, USA) and quantified by RT-qPCR in a Bio-Rad CFX384 Touch detection system using the KAPPA KK4824 kit (Kapa Biosystems, Wilmington, USA). Two 150-bp single-end sequencing libraries were prepared using the NextSeq 500/550 High Output Kit (Illumina, San Diego, USA) and sequenced on a NextSeq 500 (Illumina, San Diego, USA).

After checking sequencing quality with FastQC^141^, we used Stacks software version 1.42^142^ for demultiplexing and checking the amount of data generated for each sample. The TASSEL4-POLY pipeline^143^, developed from TASSEL-GBS^144^, was used for variant calling. Most parameters were set at their standard values; exceptions were the use of the “inclGaps” argument in the “DiscoverySNPCaller” plugin, the “misMat” argument with a value of 0.3 and the “callHets” argument in the “MergeDuplicateSNPs” plugin. Rather than aligning raw reads to a reference genome, the TASSEL-GBS pipeline first generates “tags” – unique sequences representing redundant reads – to reduce computation time^144^. We tested mapping tags against nine genomic references using two aligners: BWA version 0.7.2^145^ and Bowtie2 version 2.2.5^146^. The genomic references used were as follows: the *S. bicolor* genome^147^, the methyl-filtered genome of the sugarcane cultivar SP70-1143^47^, a sugarcane RNA-Seq assembly^148^, a de novo assembly generated from GBS data following the GBS-SNP-CROP pipeline^149^, a draft genome of the sugarcane cultivar SP80-3280^150^, a sugarcane transcriptome generated by Iso-Seq^151^, the mosaic monoploid genome of the sugarcane cultivar R570^152^, the *S. spontaneum* genome^49^ and a monoploid chromosomic set obtained from this same reference that included the “A” haplotype and unassembled scaffolds. To avoid sampling of duplicated regions, we did not include tags with multiple alignments in the ensuing analyses. After variant calling, VCFtools version 0.1.13^153^ was used to retain biallelic markers with an MAF of 0.1, no missing data and a minimum sequencing depth of 50 reads. The most appropriate reference was chosen, and adopting the method proposed by Yang et al.^43^, the ratio between alleles (allele proportions, APs) of each variant was transformed into genotypes with a fixed ploidy of 12 using the vcfR R package^154^.

### 4.5 Linkage Disequilibrium and Population Structure Analyses

For SNPs and indels, we measured LD on the ldsep R package^162^ by calculating the squared correlation coefficient (r^2^) between pairs of markers on the same chromosome. The decay of LD over physical distance was investigated by pooling all chromosomes, plotting pairwise r^2^ values against the distance between markers and fitting a curve using the equation proposed by Hill and Weir^163^. The critical r^2^ for LD decay was set to 0.1, the most commonly used threshold for determining the existence of LD^164^. Only comparisons with p < 0.05 were used in this analysis.

Three procedures were used to evaluate genetic structuring in the panel, employing dominant and codominant markers separately; for all analyses, the maximum number of clusters in the panel was set to 10. The first method was a DAPC, performed in the adegenet R package^155^. The second was PCA followed by K-means, for which missing data were imputed with the pcaMethods package^156^ and for which the optimal number of clusters was evaluated using the elbow, silhouette and gap statistic methods in the factoextra package. The last was a Bayesian clustering of genotypes into predetermined numbers of clusters (K) performed on STRUCTURE software^157^, assuming an admixture model with correlated allelic frequencies between populations. Ten independent runs were implemented for each K, and for dominant markers, estimates of probabilities of values of K in each run were taken following 100,000 generations as burn-in and 200,000 generations sampled in a Monte Carlo Markov Chain (MCMC). For Bayesian clustering using SNPs and indels, we used a subset of 7,000 markers randomly sampled from the total dataset, parallelized STRUCTURE with StrAuto software^158^ and sampled 100,000 generations in the MCMC. In both cases, the most likely number of genetic clusters was determined by the ad hoc statistics ΔK^159^ and the LnP(D) probability logarithm; the output was interpreted in STRUCTURE HARVESTER software version 0.6.94^160^. Clumpak software^161^ was used to average the admixture proportions of runs and to estimate cluster membership coefficients for genotypes.

### 4.6 Association Analyses

#### 4.6.1 FarmCPU

Association analyses with dominant markers were performed with the FarmCPU^165^ method in R For these analyses, markers were recoded to indicate the presence (0) and absence (2) of bands. We tested FarmCPU using no covariates and including matrices obtained from the three genetic structure analyses described in the previous section as such. In each case, a Q-Q)plot of the −log_10_(p) values of markers was generated, and the genomic inflation factor λ^166^ was calculated. The average λ from analyses employing each covariate matrix was calculated and used to select the model that best controlled inflation. The Bonferroni correction with α = 0.05 was used to establish the significance threshold for associations, and the phenotypic variance explained by each marker was estimated for significant marker-trait associations using a linear model in R software.

#### 4.6.2 Mixed Modeling in GWASpoly

Association analyses using SNPs and indels were performed using mixed linear model approaches in the GWASpoly R package^63^. The output of the three genetic structure analyses previously described was used to build Q matrices, which were included in the models as fixed effects. Similarly, three different genetic kinship matrices (K) of the panel were computed and included as random effects: (I) a MM^T^ matrix^167^, built on GWASpoly; (II) a complete autopolyploid matrix based on Slater et al.^168^, built with the AGHmatrix R package^169^; and (III) a pseudodiploid matrix based on Slater et al. ^168^, also built with AGHmatrix. We tested twelve Q + K combinations, and for each of them, six marker-effect models were used: general, additive, simplex dominant reference, simplex dominant alternative, diploidized general and diploidized additive. For each model, a Q-Q plot of the -log_10_(p) values of markers was generated, and λ was calculated. The average λ of all traits and models employing each Q + K combination was calculated and used to select the best set of matrices. Once this combination was chosen, Manhattan plots were generated for all models and traits. The Bonferroni and FDR correction methods with α = 0.05 were assessed to establish the significance threshold for associations.

#### 4.6.3 Machine Learning Coupled with Feature Selection

Finally, we assessed the capacity of ML strategies to predict the attribution of genotypes to the phenotypic groups identified in the HCPC analyses based on all markers, following the genomic prediction approach proposed by Aono et al.^62^. For this approach, we selected accessions successfully genotyped with both SNPs/indels and AFLPs/SSRs; missing data in dominant markers were imputed as the means. We evaluated the accuracy of eight ML algorithms: adaptive boosting (AB)^170^, decision tree (DT)^171^, Gaussian naive Bayes (GNB)^172^, Gaussian process (GP)^173^, K-nearest neighbor (KNN)^174^, MLP^175^, random forest (RF)^176^ and support vector machine (SVM)^177^, all implemented in the scikit-learn Python 3 module^178^. As a cross-validation strategy, we used a stratified K-fold (k=5) repeated 100 times for different data configurations.

We then tested five FS techniques to obtain feature importance and create subsets of marker data: gradient tree boosting (FS1)^179^, L1-based FS through a linear support vector classification system (FS2)^177^, extremely randomized trees (FS3)^180^, univariate FS using ANOVA (FS4) and RF (FS5)^176^. All FS approaches were implemented in the scikit-learn Python 3 module. We tested the differences in the accuracy between the selected FS methods using ANOVAs and multiple comparisons by Tukey’s tests implemented in the agricolae R package^181^. We also evaluated intersections between these datasets: markers selected by at least two of the five methods (Inter1); markers selected by at least two of the three best methods (Inter2); and markers selected by all three best methods (Inter3). Finally, the area under ROC curves was calculated for the best ML-FS combination and plotted using the Matplotlib library89 with Python 3.

### 4.7 Marker Mapping and Annotation

The distribution of markers identified by all analyses along *S. spontaneum* “A” chromosomes was visualized using MapChart^182^. Markers previously associated with SCYLV resistance by QTL mapping^26,29^ and GWAS^27,30^ were also retrieved and included in the map. Finally, the sequences of associated markers were annotated by aligning SSR flanking sequences or the 2000-bp window adjacent to SNPs and indels against a database comprising CDSs of the genomes of 14 Poaceae species and *Arabidopsis thaliana*^62^. For this, BLASTn^183^ was used with an E-value of 1e-30, and the best alignment of each sequence was kept for analysis.

## Acknowledgments

We thank Aline C. L. Moraes for assistance in constructing and sequencing the GBS library and Maicon Volpin for assistance in fieldwork. This work was supported by grants from the São Paulo Research Foundation (FAPESP), the Conselho Nacional de Desenvolvimento Científico e Tecnológico (CNPQ, grant 424050/2016-1), the Coordenação de Aperfeiçoamento de Pessoal de Nível Superior (CAPES, Computational Biology Program), the Littoral Polytechnic Superior School (ESPOL) and the Secretaría Nacional de Ciencia y Tecnología (SENESYT). RJGP received an MSc fellowship from CAPES (grant 88887.177386/2018-00) and MSc and PhD fellowships from FAPESP (grants 2018/18588-8 and 2019/21682-9). AHA received a PhD fellowship from FAPESP (grant 2019/03232-6). RCVB received a PhD fellowship from PAEDEx-AUIP. CCS received a PD fellowship from FAPESP (grant 2015/24346-9).

## Author Contributions

DP, MGAL, MCG, LRP and APS conceived the project and designed the experiments. RJGP, RCVB, CCS, IAA and LRP performed phenotyping. RJGP, RCVB and AEC performed genotyping. RJGP and AHA analyzed the data and interpreted the results. RJGP wrote the manuscript. All authors read and approved the manuscript.

## Competing Interests

The author(s) declare no competing interests.

## Data Availability

The raw sequencing data used in this article have been submitted to the SRA/NCBI under BioProject PRJNA702641.

## Notes

### Competing Interest Statement

The authors have declared no competing interest.

### Summary of Updates

Changes in text organization and formatting.

