## Supplementary Material for "Genome-wide approaches for the identification of markers and genes associated with sugarcane yellow leaf virus resistance"

**Supplementary Results**

1. **Phenotypic data analyses**

97 sugarcane accessions inoculated with SCYLV were evaluated for the severity of SCYL symptoms and for viral titer estimated by relative and absolute RT-qPCR quantification in two consecutive years. We observed a skew in the distribution of symptom severity data towards the absence of symptoms, and despite normalization procedures, these data did not follow a normal distribution (p = 2.2e-16). This was not the case for SCYLV titer; both the relative (p = 0.999) and absolute (p = 0.930) quantification data were successfully normalized. Plots of the distribution of the best linear unbiased predictors (BLUPs) calculated for the three traits analyzed and of the correlations between them are depicted in Supplementary Fig. 2.

2. **Genotyping**

Genotyping the 97 accessions of the panel using 12 SSRs and 35 AFLPs generated 981 polymorphic markers (167 SRR fragments and 814 AFLP fragments). Upon filtering, four genotypes (74, 93, 96 and 97) and 319 markers were removed as they presented call rates below 90% or minor allele frequency (MAF) below 10%. Therefore, 93 genotypes and 662 dominant markers were retained for further analyses that involved them.

The sequencing of the GBS library including 94 sugarcane genotypes generated 119,117,758 raw reads, 87.88% of which presented a Q-value above 30. Genotypes 31 and 77 presented very low amounts of generated data (0.19 and 1.5 KB, respectively) in contrast with the library average (598 KB) and were therefore excluded from subsequent analyses to avoid biasing. The results from tag mapping against each genomic reference using BWA and Bowtie2 aligners can be found in Supplementary Table 6. To avoid sampling of duplicated regions – which are abundant in the sugarcane genome and could bias the calculation of allele proportions (APs) – we did not include tags with multiple alignments in the ensuing analyses. BWA presented slightly superior results in terms of uniquely mapped tags and was therefore chosen as the aligning tool. Regarding the nine genomic references tested for variant calling, the methyl-filtered genome of the cultivar SP70-1143 resulted in the largest number of isolated markers among all references, yielding 60,173 SNPs and 43,054 indels after rigorous filtering (Supplementary Table 7). However, knowing that a high-quality genome assembly can greatly improve the results of GWASs and allow the estimation of long-distance LD, we opted to perform ensuing analyses with markers obtained from a reference that not only yielded a large number of markers but also provided reliable information on their position. Thus, we chose to use the monoploid chromosome set isolated from the *S. spontaneum* genome as a reference; despite not yielding the highest number of markers (38,710 SNPs and 32,178 indels), this genome had a performance superior to most of the other references tested here and, unlike many others, provided markers with information of position at chromosome level.

### 3. Genetic structure analyses

The genetic structure of the panel was investigated using two separate sets of markers – AFLPs and SSRs scored as dominant and codominant SNPs and indels with AP information – and three different approaches – a discriminant analysis on principal components (DAPC), a principal component analysis (PCA) followed by k-means and a Bayesian clustering implemented in STRUCTURE software. These three methods were first performed with dominant markers. The DAPC was performed retaining 70 principal components (PCs) that together explained ~90% of the observed variance (Supplementary Fig. 6a); the Bayesian information criterion (BIC) curve indicated k = 3 as the most likely number of clusters in the panel (Supplementary Fig. 6b). Therefore, genotypes were separated into three clusters (A1-3); the plotting of the panel elements onto the first two linear discriminants from this analysis is shown in Fig. 3a. Despite resulting in the subdivision of the panel into four clusters (B1-4), as indicated by two out of three statistical methods performed to assess the optimal number of clusters in the panel (Supplementary Fig. 7a-c), clustering by PCA followed by k-means yielded results that shared similarities with the DAPC. Genotypes forming B1 and B4, for instance, were entirely represented by A3 and A1, respectively. Fig. 3b depicts a projection of groups identified by k-means onto the PCA's first two PCs, which accounted for ~18% of the variation in the genotypic data. Last, based on Bayesian clustering, both ΔK and LnP(D) methods clearly indicated K = 2 as the most likely number of clusters in the panel (Supplementary Fig. 8a and b, respectively). Therefore, we present here a plot of the estimated association coefficients of individuals for these two clusters (Fig. 3c), named C1 and C2. Clustering by this method was highly consistent with clustering by k-means, as C1 corresponded to the union of B1 and B2, and C2 corresponded to the union of B3 and B4.

These same three analyses were next performed with codominant markers. DAPC was performed retaining 60 PCs explaining ~90% of the observed variance (Supplementary Fig. 9a); the BIC curve indicated k = 3 as the most likely number of clusters in the panel (Supplementary Fig. 9b). Thus, the panel was separated into three groups (D1-3); the plotting of the panel elements onto the first two linear discriminants from this analysis is shown in Fig. 4a. PCA followed by k-means was executed assuming the existence of two clusters (E1 and E2), as clearly indicated by all three testing methods employed (Supplementary Fig. 10a-c). Despite indicating the subdivision of the panel into two clusters, the attribution of genotypes to these groups shared similarities with the DAPC clustering. E1 corresponded to D2, and E2 was equivalent to D1 and D3 combined. Fig. 4b depicts a projection of these groups onto the PCA's first two PCs, which accounted for ~33% of the variation in the genotypic data. Finally, based on Bayesian clustering, both ΔK and LnP(D) methods indicated K = 2 as the most likely number of clusters in the panel (Supplementary Fig. 11a and b, respectively). Thus, we present here a plot of the estimated association coefficients of individuals for these two clusters, named F1 and F2 (Fig. 4c). Clustering by this procedure was also consistent with clustering by the other two methods, as F1 corresponded to the union of D2 and D3, and F2 corresponded to D1.

**Supplementary Tables**

**Supplementary Table 1. List of genotypes in the association panel with their respective parents.**

| ID Number | Genotype | Female parent | Male Parent |
|---|---|---|---|
| 1 | US571415 | *S. spontaneum* | SP701143 |
| 2 | Cana alho | *S. officinarum* | Unknown |
| 3 | IACSP012410 | R570 | IACSP933046 |
| 4 | IACSP046007 | PO8862 | IAC912195 |
| 5 | IACSP046059 | IACSP953018 | SP775181 |
| 6 | IACSP953028 | IACSP953028 | Unknown |
| 7 | NG5712 | *S. robustum* | Unknown |
| 8 | IACSP042516 | IAC912288 | CT961414 |
| 9 | Krakatau | *S. spontaneum* | Unknown |
| 10 | IN8482 | *S. spontaneum* | SP823530 |
| 11 | IACSP042508 | SP913059 | Unknown |
| 12 | IACSP955037 | SP842066 | IACSP973391 |
| 13 | IACCTC059630 | IACSP972053 | SP80185 |
| 14 | IACSP046053 | IACSP956114 | IAC873396 |
| 15 | IACSP993009 | CP701133 | Unknown |
| 16 | IS76155 | *S. officinarum* | RB855113 |
| 17 | IACSP012421 | RB835486 | SP847017 |
| 18 | IACSP042521 | IAC912288 | SP88721 |
| 19 | IACSP046058 | IACSP953018 | SP87365 |
| 20 | IACSP991020 | SP851682 | Unknown |
| 21 | IACSP045065 | IACSP956114 | RB72454 |
| 22 | IACCTC069767 | IACSP963069 | SP803280 |
| 23 | IACSP985046 | SP813251 | Unknown |
| 24 | IACSP018034 | IAC913093 | Unknown |
| 25 | IACSP993357 | SP921584 | Unknown |
| 26 | IACCTC059578 | Whitetransparent | SP963092 |
| 27 | IACCTC055580 | CTC4 | IACSP933046 |
| 28 | IACSP976628 | RB835486 | SP931124 |
| 29 | IACSP976680 | SP842066 | Unknown |
| 30 | IACCTC059534 | IACSP953028 | Unknown |
| 31 | IACSP972084 | RB855035 | SP775181 |
| 32 | CTC15 | SP842025 | Unknown |
| 33 | IAC914168 | SP813137 | IAC873396 |
| 34 | IACBIO232 | RB855465 | CTC9 |
| 35 | IACBIO241 | RB855465 | SP963092 |
| 36 | IACBIO257 | NG26011 | IAC863154 |
| 37 | IACBIO266 | NG26011 | RB855156 |
| 38 | IACBIO270 | SES069 | Unknown |
| 39 | IACBIO271 | SES069 | SP80185 |

| | | | |
|---|---|---|---|
| 40 | IACBIO273 | SES069 | Unknown |
| 41 | IACBIO275 | SES069 | Unknown |
| 42 | IACBIO277 | SES069 | IACSP972077 |
| 43 | IACBIO279 | SES069 | IAC873396 |
| 44 | IACCTC053616 | IAC915155 | RB85453 |
| 45 | IACCTC056518 | SP911049 | SP832847 |
| 46 | IACCTC059552 | IACSP955011 | SP775158 |
| 47 | IACCTC059607 | IACSP972053 | SP963092 |
| 48 | IACCTC059634 | IACSP972053 | IACSP953050 |
| 49 | IACCTC061050 | CT943165 | IACSP952078 |
| 50 | IACCTC069708 | IACSP962042 | SP823697 |
| 51 | IACCTC069713 | IACSP955000 | SP87432 |
| 52 | IACCTC069741 | IACSP955000 | Unknown |
| 53 | IACSP012417 | RB835486 | RB835486 |
| 54 | IACSP012430 | IAC912195 | CT961414 |
| 55 | IACSP015501 | SP913059 | SP80185 |
| 56 | IACSP015519 | RB835486 | Unknown |
| 57 | IACSP018046 | R570 | Glagah |
| 58 | IACSP018082 | IAC913093 | Glagah |
| 59 | IACSP018158 | RB835486 | Unknown |
| 60 | IACSP022067 | SP973060 | Unknown |
| 61 | IACSP022125 | SP901161 | Unknown |
| 62 | IACSP023025 | SP924230 | Unknown |
| 63 | IACSP023168 | IAC914216 | Unknown |
| 64 | IACSP042503 | PO8862 | Unknown |
| 65 | IACSP042504 | IACSP955037 | Unknown |
| 66 | IACSP042509 | SP913059 | CT961414 |
| 67 | IACSP042510 | IAC913093 | CT931231 |
| 68 | IACSP043123 | IACSP953028 | RB855453 |
| 69 | IACSP043148 | IACSP953018 | IAC911099 |
| 70 | IACSP043150 | IACSP953018 | CT014455 |
| 71 | IACSP043259 | IACSP966026 | CT961307 |
| 72 | IACSP045081 | IACSP966026 | SP775181 |
| 73 | IACSP046032 | IACSP953028 | CTC957 |
| 74 | IACSP046035 | SP913059 | SP775181 |
| 75 | IACSP046073 | IACSP953028 | SP701143 |
| 76 | IACSP046077 | IACSP972109 | SP901644 |
| 77 | IACSP046152 | IAC911099 | SP88813 |
| 78 | IACSP933046 | SP791011 | SP803280 |
| 79 | IACSP953018 | SP842189 | CTC9 |
| 80 | IACSP956114 | IAC873187 | CTC9638 |
| 81 | IACSP962008 | SP80144 | SP832847 |
| 82 | IACSP963056 | SP826108 | IACSP933046 |

| | | | |
|---|---|---|---|
| 83 | IACSP963069 | SP803280 | IACSP933046 |
| 84 | IACSP963076 | SP847017 | RB855453 |
| 85 | IACSP973384 | RB855113 | IACSP966026 |
| 86 | IACSP974039 | RB835486 | SP924221 |
| 87 | IACSP974048 | SP842066 | IACSP956114 |
| 88 | IACSP982053 | SP847017 | SP801842 |
| 89 | IACSP983011 | IAC913093 | CTC9019 |
| 90 | IACSP993369 | SP901616 | IACCTC059607 |
| 91 | IACSP994011 | SP842025 | RB855453 |
| 92 | IJ76293 | *S. robustum* | IAC863154 |
| 93 | IN8458 | *S. spontaneum* | SP801842 |
| 94 | IN8488 | *S. spontaneum* | Unknown |
| 95 | NG57213 | Unknown | Unknown |
| 96 | RB935744 | RB835089 | TUC717 |
| 97 | SP832847 | HJ5741 | Unknown |

**Supplementary Table 2. Combinations of primers used for amplified fragment length polymorphism (AFLP) genotyping.** Sequences correspond to the three selective nucleotides located at the 3' end of *Eco*RI and *Msp*I primers.

| **Combination ID** | **3' Primer sequence** | |
|---|---|---|
| | ***Eco*RI** | ***Msp*I** |
| 1 | AGA-IRD800 | ACA |
| 2 | AGA-IRD800 | TTG |
| 3 | AGA-IRD800 | ACC |
| 5 | AGA-IRD800 | TAG |
| 6 | AGA-IRD800 | CGC |
| 9 | AGA-IRD800 | GAA |
| 10 | ACA-IRD700 | ACA |
| 11 | ACA-IRD700 | TTG |
| 12 | ACA-IRD700 | ACC |
| 14 | ACA-IRD700 | TAG |
| 15 | ACA-IRD700 | CGC |
| 16 | ACA-IRD700 | GAG |
| 18 | ACA-IRD700 | GAA |
| 20 | ACC-IRD700 | TTG |
| 21 | ACC-IRD700 | ACC |
| 22 | ACC-IRD700 | TCG |
| 24 | ACC-IRD700 | CGC |
| 25 | ACC-IRD700 | GAG |
| 26 | ACC-IRD700 | ACT |
| 33 | AAC-IRD700 | CGC |
| 34 | AAC-IRD700 | GAG |
| 36 | AAC-IRD700 | GAA |
| 37 | ACG-IRD700 | ACA |
| 40 | ACG-IRD700 | TCG |
| 41 | ACG-IRD700 | TAG |
| 42 | ACG-IRD700 | CGC |
| 48 | AGG-IRD800 | ACC |
| 49 | AGG-IRD800 | TCG |
| 51 | AGG-IRD800 | CGC |
| 53 | AGG-IRD800 | ACT |
| 54 | AGG-IRD800 | GAA |
| 55 | AGG-IRD800 | ACA |
| 56 | AGG-IRD800 | TTG |
| 59 | AGG-IRD800 | TAG |
| 60 | AGG-IRD800 | CGC |

**Supplementary Table 3. Locus name, repetition motif, amplicon size range, annealing temperature (Ta) and sequence of the primers used to amplify the simple sequence repeat (SSR) loci used in the study.**

| SSR | Motif | Amplicon size range (bp) | Ta (°C) | Forward/Reverse primer sequences |
|---|---|---|---|---|
| CV29 | $(ATCT)_{14}$ | 85-151 | 60.0 | TCGCGTCCACCAATGTAACC/GCGTGCATCGCTTGTGTCTT |
| CV37 | $(TTTC)_{15}$ | 114-171 | 60.0 | GGATGGACGACGTGTCCTGG/ATAAAGTGGCCGCTTGGATTGA |
| CV38 | $(CTTTT)_{18}$ | 93-204 | 60.0 | GAAGCAGGGGCCTCAAGTTG/GTCAAACAGGCGATCTGGCTC |
| CV60 | $(CTCTCC)_{5}$ | 158-194 | 60.0 | AATCTGCACCCTGCCCTCTC/CAGCTGGAGCATGGATGGAG |
| CV79 | $(CTATAT)_{11}(TATAGA)_{6}$ | 131-231 | 60.0 | GGCACTGCTGGTGGTTGATTG/TCCCACATCAAGAGGCAGCTA |
| CV94 | $(AAAAAG)_{5}(CGT)_{5}$ | 182-234 | 60.0 | GGCAGGCCAAGATGAATGAAG/AGCACAGCGGAGGGTACGG |
| CV106 | $(GGC)_{8}$ | 78-162 | 60.0 | AAACAGAGCATACTCGAGGCC/ACGTTGCTGACGAGGTTTTCC |
| SCB213 | $(TCC)_{6}$ | 247-337 | 62.0 | AGCCGTCAGGGGTCAGG/ATTCGATGGAGCCTGAGTGAG |
| SCB312 | $(TCC)_{7}$ | 190-278 | 64.7 | AGTCCGTCGCCGTAATCATCTTG/GCACCTCCTCCTTTCCTTCCTTATT |
| SCB381 | $(TAC)_{8}$ | 190-257 | 60.0 | TGGAGCTCCGTCTTCTTGTT/GCTAGCCCGTACATTGGGTA |
| SCB423 | $(TTG)_{5}$ | 250-300 | 61 | CCATGTGGCTTCCTGAAACT/ACAGGCACTTCAAGGGAAGA |
| SCB436 | $(GAG)_{5}$ | 158-274 | 60.0 | AGTACGCCTGAGTCCTGACG/AGGTGCAAGGGCTGATAGAA |

**Supplementary Table 4. Distribution of the 97 panel accessions among clusters identified by two hierarchical clustering on principal component (HCPC) analyses performed with best linear unbiased predictor (BLUP) values.** Genotypes were divided into three clusters (Q1, Q2 and Q3) based on SCYLV titer determined by RT-qPCR in the first HCPC and into three clusters (SQ1, SQ2 and SQ3) based on SCYL symptom severity and SCYLV titer in the second HCPC.

| ID Number | Q Cluster | SQ Cluster | ID Number | Q Cluster | SQ Cluster | ID Number | Q Cluster | SQ Cluster |
|---|---|---|---|---|---|---|---|---|
| **1** | Q2 | SQ2 | **34** | Q2 | SQ2 | **67** | Q1 | SQ1 |
| **2** | Q2 | SQ2 | **35** | Q1 | SQ1 | **68** | Q3 | SQ2 |
| **3** | Q3 | SQ2 | **36** | Q2 | SQ1 | **69** | Q3 | SQ2 |
| **4** | Q1 | SQ1 | **37** | Q1 | SQ1 | **70** | Q3 | SQ3 |
| **5** | Q2 | SQ2 | **38** | Q1 | SQ1 | **71** | Q2 | SQ2 |
| **6** | Q3 | SQ2 | **39** | Q1 | SQ1 | **72** | Q3 | SQ2 |
| **7** | Q3 | SQ2 | **40** | Q1 | SQ1 | **73** | Q2 | SQ2 |
| **8** | Q2 | SQ2 | **41** | Q1 | SQ1 | **74** | Q2 | SQ2 |
| **9** | Q2 | SQ1 | **42** | Q3 | SQ2 | **75** | Q3 | SQ2 |
| **10** | Q2 | SQ1 | **43** | Q1 | SQ1 | **76** | Q3 | SQ2 |
| **11** | Q2 | SQ2 | **44** | Q1 | SQ1 | **77** | Q2 | SQ2 |
| **12** | Q3 | SQ2 | **45** | Q2 | SQ3 | **78** | Q3 | SQ2 |
| **13** | Q3 | SQ2 | **46** | Q3 | SQ2 | **79** | Q3 | SQ2 |
| **14** | Q3 | SQ2 | **47** | Q3 | SQ2 | **80** | Q3 | SQ2 |
| **15** | Q2 | SQ3 | **48** | Q2 | SQ2 | **81** | Q3 | SQ2 |
| **16** | Q3 | SQ2 | **49** | Q2 | SQ2 | **82** | Q2 | SQ2 |
| **17** | Q3 | SQ2 | **50** | Q3 | SQ2 | **83** | Q3 | SQ3 |
| **18** | Q3 | SQ3 | **51** | Q3 | SQ2 | **84** | Q3 | SQ2 |
| **19** | Q3 | SQ2 | **52** | Q3 | SQ2 | **85** | Q3 | SQ2 |
| **20** | Q3 | SQ2 | **53** | Q3 | SQ2 | **86** | Q3 | SQ2 |
| **21** | Q3 | SQ2 | **54** | Q2 | SQ2 | **87** | Q3 | SQ2 |
| **22** | Q3 | SQ2 | **55** | Q3 | SQ2 | **88** | Q2 | SQ2 |
| **23** | Q1 | SQ1 | **56** | Q2 | SQ2 | **89** | Q2 | SQ2 |
| **24** | Q3 | SQ2 | **57** | Q3 | SQ3 | **90** | Q3 | SQ2 |
| **25** | Q3 | SQ2 | **58** | Q2 | SQ3 | **91** | Q2 | SQ2 |
| **26** | Q2 | SQ2 | **59** | Q3 | SQ2 | **92** | Q2 | SQ1 |
| **27** | Q2 | SQ2 | **60** | Q2 | SQ2 | **93** | Q2 | SQ2 |
| **28** | Q3 | SQ2 | **61** | Q3 | SQ2 | **94** | Q1 | SQ1 |
| **29** | Q2 | SQ2 | **62** | Q2 | SQ2 | **95** | Q3 | SQ2 |
| **30** | Q2 | SQ2 | **63** | Q2 | SQ2 | **96** | Q3 | SQ2 |
| **31** | Q3 | SQ2 | **64** | Q2 | SQ2 | **97** | Q2 | SQ2 |
| **32** | Q2 | SQ2 | **65** | Q2 | SQ2 | | | |
| **33** | Q2 | SQ2 | **66** | Q2 | SQ2 | | | |

**Supplementary Table 5. P-values associated with Dunn tests comparing the best linear unbiased predictor (BLUP) values of traits between the clusters identified by the two hierarchical clustering on principal components (HCPC) analyses.** Asterisks (*) indicate statistical significance ($p < 0.05$).

| | Symptom severity | Relative quantification | Absolute quantification |
|---|---|---|---|
| **Q1 vs. Q2** | - | 0.0037* | 0.0008* |
| **Q1 vs. Q3** | - | 0.0000* | 0.0000* |
| **Q2 vs. Q3** | - | 0.0000* | 0.0000* |
| **SQ1 vs. SQ2** | 0.0003* | 0.0000* | 0.0000* |
| **SQ1 vs. SQ3** | 0.0000* | 0.0004* | 0.0056* |
| **SQ2 vs. SQ3** | 0.0001* | 1 | 0.4037 |

**Supplementary Table 6. Percentage of tag mapping against the nine chosen references using BWA and Bowtie2.**

| Reference | | BWA | | | | Bowtie2 | | | |
|---|---|---|---|---|---|---|---|---|---|
| | | Unmapped | Mapped | | | Unmapped | Mapped | | |
| | | | 1 Time | > 1 Time | Total | | 1 Time | > 1 Time | Total |
| ***S. bicolor*** **genome** | | 71.64% | 24.00% | 4.36% | 28.36% | 64.72% | 25.99% | 9.30% | 35.28% |
| **Methyl-filtered genome** | | 22.40% | 62.50% | 15.10% | 77.60% | 18.13% | 49.20% | 32.67% | 81.87% |
| **RNA-Seq** | | 75.34% | 24.04% | 0.62% | 24.66% | 72.34% | 24.87% | 2.79% | 27.66% |
| **De novo assembly** | | 0.58% | 20.78% | 78.62% | 99.40% | 0.13% | 4.10% | 95.77% | 99.87% |
| **Draft genome** | | 25.56% | 29.52% | 44.93% | 74.44% | 21.91% | 22.28% | 55.81% | 78.09% |
| **Iso-Seq** | | 79.61% | 7.93% | 12.46% | 20.39% | 62.69% | 11.02% | 26.29% | 37.31% |
| **Mosaic monoploid genome** | | 45.34% | 39.12% | 15.53% | 54.66% | 41.38% | 35.40% | 23.22% | 58.62% |
| ***S. spontaneum*** **genome** | **Complete** | 11.09% | 11.73% | 77.18% | 88.91% | 8.09% | 5.46% | 86.44% | 91.91% |
| | **Monoploid set** | 28.13% | 38.34% | 33.53% | 71.87% | 24.21% | 32.50% | 43.29% | 75.79% |

**Supplementary Table 7. Number of single-nucleotide polymorphisms (SNPs) and insertion and deletion (indel) markers isolated by mapping sequencing tags to each genomic reference.**

| Reference | | Prefiltering | | | Postfiltering | | |
|---|---|---|---|---|---|---|---|
| | | **Total markers** | **SNPs** | **Indels** | **Total markers** | **SNPs** | **Indels** |
| ***S. bicolor* genome** | | 451,845 | 288,387 | 163,458 | 49,053 | 28,674 | 20,379 |
| **Methyl-filtered genome** | | 1,336,253 | 870,075 | 466,178 | 103,096 | 60,171 | 42,925 |
| **RNA-Seq** | | 417,588 | 292,943 | 124,645 | 43,739 | 28,934 | 14,805 |
| **De novo assembly** | | 120,419 | 120,419 | 21,274 | 133 | 123 | 10 |
| **Draft Genome** | | 681,836 | 443,727 | 238,109 | 41,169 | 23,568 | 17,601 |
| **Iso-Seq** | | 151,783 | 105,796 | 45,987 | 13,147 | 8,479 | 4,668 |
| **Mosaic monoploid genome** | | 882,838 | 564,585 | 318,253 | 74,473 | 42,750 | 31,723 |
| ***S. spontaneum* genome** | **Complete** | 294,525 | 192,548 | 101,977 | 6,184 | 3,625 | 2,559 |
| | **Monoploid set** | 940,255 | 574,962 | 365,293 | 70,978 | 38,755 | 32,223 |

**Supplementary Table 8. Allocation of panel genotypes to clusters identified by population structure analyses.** Two separate sets of markers (dominant and codominant) and three clustering methods were employed: a discriminant analysis of principal components (DAPC), a principal component analysis (PCA) followed by k-means and a Bayesian clustering analysis.

| ID Number | Dominant DAPC | Dominant k-means | Dominant Bayesian | Codominant DAPC | Codominant k-means | Codominant Bayesian |
|---|---|---|---|---|---|---|
| 1 | A3 | B2 | C1 | D2 | E1 | F1 |
| 2 | A3 | B2 | C1 | D2 | E1 | F1 |
| 3 | A3 | B2 | C1 | D3 | E2 | F1 |
| 4 | A3 | B2 | C1 | D2 | E1 | F1 |
| 5 | A3 | B2 | C1 | D2 | E1 | F1 |
| 6 | A3 | B1 | C1 | D2 | E1 | F1 |
| 7 | A3 | B1 | C1 | D2 | E1 | F1 |
| 8 | A3 | B2 | C1 | D2 | E1 | F1 |
| 9 | A1 | B3 | C2 | D2 | E1 | F1 |
| 10 | A1 | B3 | C2 | D2 | E1 | F1 |
| 11 | A3 | B2 | C1 | D2 | E1 | F1 |
| 12 | A3 | B2 | C1 | D2 | E1 | F1 |
| 13 | A3 | B2 | C1 | D2 | E1 | F1 |
| 14 | A3 | B1 | C1 | D2 | E1 | F1 |
| 15 | A3 | B2 | C1 | D2 | E1 | F1 |
| 16 | A3 | B2 | C1 | D2 | E1 | F1 |
| 17 | A3 | B2 | C1 | D1 | E2 | F2 |
| 18 | A3 | B2 | C1 | D2 | E1 | F1 |
| 19 | A3 | B2 | C1 | D2 | E1 | F1 |
| 20 | A3 | B2 | C1 | D2 | E1 | F1 |
| 21 | A3 | B2 | C1 | D2 | E1 | F1 |
| 22 | A3 | B2 | C1 | D2 | E1 | F1 |
| 23 | A3 | B2 | C1 | D2 | E1 | F1 |
| 24 | A3 | B2 | C1 | D2 | E1 | F1 |
| 25 | A3 | B2 | C1 | D1 | E2 | F2 |
| 26 | A3 | B2 | C1 | D2 | E1 | F1 |
| 27 | A3 | B2 | C1 | D2 | E1 | F1 |
| 28 | A3 | B1 | C1 | D3 | E2 | F1 |
| 29 | A3 | B2 | C1 | D1 | E2 | F2 |
| 30 | A3 | B1 | C1 | - | - | - |
| 31 | A3 | B2 | C1 | D2 | E1 | F1 |
| 32 | A3 | B2 | C1 | D2 | E1 | F1 |
| 33 | A3 | B2 | C1 | D2 | E1 | F1 |
| 34 | A3 | B2 | C1 | D2 | E1 | F1 |
| 35 | A3 | B2 | C1 | D1 | E2 | F2 |

| | | | | | | |
|---|---|---|---|---|---|---|
| 36 | A3 | B2 | C1 | D2 | E1 | F1 |
| 37 | A3 | B2 | C1 | D2 | E1 | F1 |
| 38 | A1 | B3 | C2 | D2 | E1 | F1 |
| 39 | A1 | B3 | C2 | D2 | E1 | F1 |
| 40 | A1 | B4 | C2 | D3 | E2 | F1 |
| 41 | A1 | B3 | C2 | D1 | E2 | F2 |
| 42 | A1 | B4 | C2 | D2 | E1 | F1 |
| 43 | A1 | B3 | C2 | D2 | E1 | F1 |
| 44 | A3 | B2 | C1 | D2 | E1 | F1 |
| 45 | A3 | B2 | C1 | D2 | E1 | F1 |
| 46 | A3 | B2 | C1 | D2 | E1 | F1 |
| 47 | A3 | B2 | C1 | D1 | E2 | F2 |
| 48 | A3 | B2 | C1 | D2 | E1 | F1 |
| 49 | A3 | B2 | C1 | D2 | E1 | F1 |
| 50 | A3 | B2 | C1 | D2 | E1 | F1 |
| 51 | A3 | B2 | C1 | D2 | E1 | F1 |
| 52 | A3 | B1 | C1 | D3 | E2 | F1 |
| 53 | A3 | B2 | C1 | D2 | E1 | F1 |
| 54 | A3 | B2 | C1 | D2 | E1 | F1 |
| 55 | A3 | B2 | C1 | D2 | E1 | F1 |
| 56 | A3 | B2 | C1 | D2 | E1 | F1 |
| 57 | A3 | B2 | C1 | D2 | E1 | F1 |
| 58 | A3 | B1 | C1 | D2 | E1 | F1 |
| 59 | A3 | B2 | C1 | D1 | E2 | F2 |
| 60 | A3 | B1 | C1 | D2 | E1 | F1 |
| 61 | A3 | B2 | C1 | D2 | E1 | F1 |
| 62 | A3 | B2 | C1 | D2 | E1 | F1 |
| 63 | A3 | B2 | C1 | D2 | E1 | F1 |
| 64 | A2 | B3 | C2 | D3 | E2 | F1 |
| 65 | A2 | B2 | C1 | D2 | E1 | F1 |
| 66 | A2 | B3 | C2 | D2 | E1 | F1 |
| 67 | A2 | B2 | C1 | D2 | E1 | F1 |
| 68 | A2 | B2 | C1 | D2 | E1 | F1 |
| 69 | A2 | B3 | C2 | D2 | E1 | F1 |
| 70 | A2 | B3 | C2 | D2 | E1 | F1 |
| 71 | A2 | B3 | C2 | D2 | E1 | F1 |
| 72 | A2 | B2 | C1 | D2 | E1 | F1 |
| 73 | A2 | B3 | C2 | D2 | E1 | F1 |
| 74 | - | - | - | D2 | E1 | F1 |
| 75 | A2 | B3 | C2 | D2 | E1 | F1 |
| 76 | A2 | B3 | C2 | - | - | - |
| 77 | A2 | B3 | C2 | D2 | E1 | F1 |
| 78 | A2 | B3 | C2 | D2 | E1 | F1 |

| | | | | | | |
|---|---|---|---|---|---|---|
| 79 | A2 | B3 | C2 | D2 | E1 | F1 |
| 80 | A2 | B2 | C1 | D2 | E1 | F1 |
| 81 | A2 | B3 | C2 | D2 | E1 | F1 |
| 82 | A2 | B2 | C1 | D2 | E1 | F1 |
| 83 | A2 | B2 | C1 | D2 | E1 | F1 |
| 84 | A2 | B3 | C2 | D2 | E1 | F1 |
| 85 | A2 | B3 | C2 | D2 | E1 | F1 |
| 86 | A2 | B3 | C2 | D2 | E1 | F1 |
| 87 | A2 | B2 | C1 | - | - | - |
| 88 | A2 | B3 | C2 | - | - | - |
| 89 | A2 | B3 | C2 | D2 | E1 | F1 |
| 90 | A2 | B3 | C2 | D1 | E2 | F2 |
| 91 | A2 | B3 | C2 | D2 | E1 | F1 |
| 92 | A1 | B3 | C2 | D2 | E1 | F1 |
| 93 | - | - | - | D2 | E1 | F1 |
| 94 | A1 | B3 | C2 | D2 | E1 | F1 |
| 95 | A2 | B3 | C2 | - | - | - |
| 96 | - | - | - | D2 | E1 | F1 |
| 97 | - | - | - | D2 | E1 | F1 |

**Supplementary Table 9. Dominant markers significantly associated with the three analyzed traits using the fixed and random model circulating probability unification (FarmCPU) method.** For each marker-trait association, the trait, $-\log_{10}(p)$ value and the phenotypic variance explained by each marker (adjusted $r^2$) are provided.

| Marker | Trait | $-\log_{10}(p)$ | Effect | $r^2$ |
|---|---|---|---|---|
| MSP_ACT-ECO_AGG-800-0106 | Symptom severity | 2.79 | -1.57 | 0.116 |
| MSP_TCG-ECO_ACG_700-0240 | Relative quantification | 2.55 | -0.14 | 0.234 |
| MSP_TCG-ECO_ACG_700-0268 | Relative quantification | 2.86 | 0.15 | 0.249 |
| MSP_TTG-ECO_ACA-700-0133 | Relative quantification | 2.85 | 0.19 | 0.244 |
| MSP_TTG-ECO_ACA-700-0238 | Relative quantification | 8.42 | 0.27 | 0.279 |
| MSP_TTG-ECO_AGA-800-0100 | Relative quantification | 2.61 | 0.13 | 0.213 |
| MSP_ACT-ECO_AGG-800-0116 | Absolute quantification | 9.09 | 0.44 | 0.302 |
| MSP_CGC-ECO_AGG-800-0148 | Absolute quantification | 4.23 | 0.21 | 0.251 |
| MSP_GAA-ECO_AGG-800-0207 | Absolute quantification | 2.4 | 0.15 | 0.099 |
| MSP_TTG-ECO_AAG_800_2-0133 | Absolute quantification | 2.7 | 0.25 | 0.280 |
| MSP_TTG-ECO_ACA-700-0133 | Absolute quantification | 3.8 | 0.32 | 0.222 |

**Supplementary Table 10. Codominant markers significantly associated with the three analyzed traits.** For each marker-trait association, the chromosome location, position number, trait, marker-effect model used, $-\log_{10}(p)$ value, effect and polymorphism (reference/alternative alleles) are provided. Effects are not provided for the general and diploidized general models.

| Marker | Chromosome | Position | Trait | Model | $-\log_{10}(p)$ | Effect | Polymorphism |
|---|---|---|---|---|---|---|---|
| S1_3322004 | 1 | 3322004 | Symptom severity | General | 6.97 | NA | C/T |
| S1_7029466 | 1 | 7029466 | Symptom severity | Diplo-general | 6.71 | NA | T/C |
| S1_7029466 | 1 | 7029466 | Symptom severity | Diplo-additive | 6.71 | -8.18 | T/C |
| S1_7029466 | 1 | 7029466 | Symptom severity | 1-dom-alt | 6.71 | -8.18 | T/C |
| S1_7029472 | 1 | 7029472 | Symptom severity | Diplo-general | 6.71 | NA | -/C |
| S1_7029472 | 1 | 7029472 | Symptom severity | Diplo-additive | 6.71 | -8.18 | -/C |
| S1_7029472 | 1 | 7029472 | Symptom severity | 1-dom-alt | 6.71 | -8.18 | -/C |
| S1_7029473 | 1 | 7029473 | Symptom severity | Diplo-general | 6.71 | NA | -/C |
| S1_7029473 | 1 | 7029473 | Symptom severity | Diplo-additive | 6.71 | -8.18 | -/C |
| S1_7029473 | 1 | 7029473 | Symptom severity | 1-dom-alt | 6.71 | -8.18 | -/C |
| S1_7029474 | 1 | 7029474 | Symptom severity | Diplo-general | 6.71 | NA | -/C |
| S1_7029474 | 1 | 7029474 | Symptom severity | Diplo-additive | 6.71 | -8.18 | -/C |
| S1_7029474 | 1 | 7029474 | Symptom severity | 1-dom-alt | 6.71 | -8.18 | -/C |
| S1_7029475 | 1 | 7029475 | Symptom severity | Diplo-general | 6.71 | NA | -/C |
| S1_7029475 | 1 | 7029475 | Symptom severity | Diplo-additive | 6.71 | -8.18 | -/C |
| S1_7029475 | 1 | 7029475 | Symptom severity | 1-dom-alt | 6.71 | -8.18 | -/C |
| S1_7029479 | 1 | 7029479 | Symptom severity | Diplo-general | 6.71 | NA | C/- |
| S1_7029479 | 1 | 7029479 | Symptom severity | Diplo-additive | 6.71 | -8.18 | C/- |
| S1_7029479 | 1 | 7029479 | Symptom severity | 1-dom-alt | 6.71 | -8.18 | C/- |
| S1_7029480 | 1 | 7029480 | Symptom severity | Diplo-general | 6.71 | NA | C/- |
| S1_7029480 | 1 | 7029480 | Symptom severity | Diplo-additive | 6.71 | -8.18 | C/- |
| S1_7029480 | 1 | 7029480 | Symptom severity | 1-dom-alt | 6.71 | -8.18 | C/- |
| S1_7029481 | 1 | 7029481 | Symptom severity | Diplo-general | 6.71 | NA | C/- |
| S1_7029481 | 1 | 7029481 | Symptom severity | Diplo-additive | 6.71 | -8.18 | C/- |
| S1_7029481 | 1 | 7029481 | Symptom severity | 1-dom-alt | 6.71 | -8.18 | C/- |

| | | | | | | | |
|---|---|---|---|---|---|---|---|
| S1_7029482 | 1 | 7029482 | Symptom severity | Diplo-general | 6.71 | NA | A/- |
| S1_7029482 | 1 | 7029482 | Symptom severity | Diplo-additive | 6.71 | -8.18 | A/- |
| S1_7029482 | 1 | 7029482 | Symptom severity | 1-dom-alt | 6.71 | -8.18 | A/- |
| S1_7029483 | 1 | 7029483 | Symptom severity | Diplo-general | 6.71 | NA | T/- |
| S1_7029483 | 1 | 7029483 | Symptom severity | Diplo-additive | 6.71 | -8.18 | T/- |
| S1_7029483 | 1 | 7029483 | Symptom severity | 1-dom-alt | 6.71 | -8.18 | T/- |
| S1_19840994 | 1 | 19840994 | Symptom severity | General | 6.11 | NA | A/G |
| S1_77494328 | 1 | 77494328 | Symptom severity | General | 6.42 | NA | A/- |
| S1_77494331 | 1 | 77494331 | Symptom severity | General | 6.42 | NA | G/C |
| S2_80240870 | 2 | 80240870 | Symptom severity | General | 6.75 | NA | C/T |
| S2_80240899 | 2 | 80240899 | Symptom severity | General | 7.06 | NA | C/T |
| S2_80240911 | 2 | 80240911 | Symptom severity | General | 6.73 | NA | G/C |
| S2_80240919 | 2 | 80240919 | Symptom severity | General | 6.76 | NA | T/- |
| S2_80240920 | 2 | 80240920 | Symptom severity | General | 6.93 | NA | A/T |
| S3_12947458 | 3 | 12947458 | Symptom severity | General | 8.74 | NA | T/C |
| S3_48163530 | 3 | 48163530 | Symptom severity | General | 7.06 | NA | T/C |
| S3_54754895 | 3 | 54754895 | Symptom severity | General | 6.14 | NA | A/G |
| S4_11629587 | 4 | 11629587 | Symptom severity | General | 6.42 | NA | T/C |
| S4_58593938 | 4 | 58593938 | Symptom severity | General | 7 | NA | G/A |
| S4_62622513 | 4 | 62622513 | Symptom severity | Diplo-general | 6.11 | NA | T/- |
| S4_62622513 | 4 | 62622513 | Symptom severity | Diplo-additive | 6.11 | -5.21 | T/- |
| S4_62622513 | 4 | 62622513 | Symptom severity | 1-dom-alt | 6.11 | -5.21 | T/- |
| S5_78687022 | 5 | 78687022 | Symptom severity | General | 7.53 | NA | C/G |
| S6_19432542 | 6 | 19432542 | Symptom severity | General | 7.41 | NA | A/G |
| S6_20347039 | 6 | 20347039 | Symptom severity | General | 6.53 | NA | -/A |
| S6_58198630 | 6 | 58198630 | Symptom severity | General | 7.09 | NA | A/T |
| S8_10096401 | 8 | 10096401 | Symptom severity | General | 6.88 | NA | C/- |
| S8_10096402 | 8 | 10096402 | Symptom severity | General | 6.88 | NA | T/- |
| S8_10096403 | 8 | 10096403 | Symptom severity | General | 6.88 | NA | T/- |
| S8_10096406 | 8 | 10096406 | Symptom severity | General | 6.88 | NA | T/A |
| S8_10096428 | 8 | 10096428 | Symptom severity | General | 6.23 | NA | T/G |

| | | | | | | | |
|---|---|---|---|---|---|---|---|
| SS_190207740 | - | 190207740 | Symptom severity | General | 6.88 | NA | A/G |
| S1_11503096 | 1 | 11503096 | Relative quantification | Diplo-general | 4.05 | NA | -/T |
| S1_11503096 | 1 | 11503096 | Relative quantification | Diplo-additive | 4.82 | -0.96 | -/T |
| S1_11503097 | 1 | 11503097 | Relative quantification | Diplo-general | 4.05 | NA | -/T |
| S1_11503097 | 1 | 11503097 | Relative quantification | Diplo-additive | 4.82 | -0.96 | -/T |
| S1_11503112 | 1 | 11503112 | Relative quantification | Diplo-general | 4.05 | NA | T/- |
| S1_11503112 | 1 | 11503112 | Relative quantification | Diplo-additive | 4.82 | -0.96 | T/- |
| S1_11503113 | 1 | 11503113 | Relative quantification | Diplo-general | 4.05 | NA | T/- |
| S1_11503113 | 1 | 11503113 | Relative quantification | Diplo-additive | 4.82 | -0.96 | A/- |
| S1_105811826 | 1 | 105811826 | Relative quantification | Diplo-general | 4.17 | NA | A/G |
| S1_105811826 | 1 | 105811826 | Relative quantification | Diplo-additive | 4.17 | 0.68 | A/G |
| S1_105811826 | 1 | 105811826 | Relative quantification | 1-dom-alt | 4.17 | 0.68 | A/G |
| S2_100207393 | 2 | 100207393 | Relative quantification | General | 4.16 | NA | G/C |
| S2_101252947 | 2 | 101252947 | Relative quantification | General | 4.67 | NA | -/A |
| S2_111395374 | 2 | 111395374 | Relative quantification | Additive | 4.61 | -0.17 | C/T |
| S3_5029059 | 3 | 5029059 | Relative quantification | General | 4.07 | NA | A/- |
| S3_5029059 | 3 | 5029059 | Relative quantification | Additive | 4.17 | -0.19 | A/- |
| S3_38170436 | 3 | 38170436 | Relative quantification | General | 4.38 | NA | C/A |
| S3_51566876 | 3 | 51566876 | Relative quantification | General | 4.34 | NA | G/- |
| S3_51566877 | 3 | 51566877 | Relative quantification | General | 4.34 | NA | C/- |
| S3_51566897 | 3 | 51566897 | Relative quantification | General | 4.44 | NA | T/C |
| S3_51566905 | 3 | 51566905 | Relative quantification | General | 4.44 | NA | T/C |
| S3_51566929 | 3 | 51566929 | Relative quantification | General | 5 | NA | A/C |
| S8_2650197 | 8 | 2650197 | Relative quantification | General | 4 | NA | -/A |
| SS_100753956 | - | 100753956 | Relative quantification | Additive | 4.49 | -0.1 | C/T |
| S2_3440535 | 2 | 3440535 | Absolute quantification | General | 4.88 | NA | A/T |
| S2_69603253 | 2 | 69603253 | Absolute quantification | General | 4.25 | NA | A/T |
| S2_69603262 | 2 | 69603262 | Absolute quantification | General | 4.01 | NA | A/C |
| S2_69603265 | 2 | 69603265 | Absolute quantification | General | 4.31 | NA | T/A |
| S2_69603295 | 2 | 69603295 | Absolute quantification | General | 4.31 | NA | A/T |
| S2_121155445 | 2 | 121155445 | Absolute quantification | General | 4.2 | NA | A/- |

| | | | | | | | |
|---|---|---|---|---|---|---|---|
| S3_5029059 | 3 | 5029059 | Absolute quantification | General | 4.19 | NA | A/- |
| S7_25132085 | 7 | 25132085 | Absolute quantification | Diplo-general | 4.37 | NA | C/T |
| S7_25132085 | 7 | 25132085 | Absolute quantification | Diplo-additive | 4.37 | -1.4 | C/T |
| S7_25132085 | 7 | 25132085 | Absolute quantification | 1-dom-alt | 4.37 | -1.4 | C/T |
| S7_76514321 | 7 | 76514321 | Absolute quantification | General | 4.32 | NA | C/- |

**Supplementary Table 11. Cross-validation results of the predictive accuracy of each machine learning (ML) model and feature selection (FS) combination employed to predict clustering by SCYLV titer determined by RT-qPCR (Q).** The machine learning models tested were adaptive boosting (AB), decision tree (DT), Gaussian naive Bayes (GNB), Gaussian process (GP), K-nearest neighbor (KNN), multilayer perceptron (MLP), random forest (RF) and support vector machine (SVM). FS methods tested were gradient tree boosting (FS1), L1-based FS through a linear support vector classification system (FS2), extremely randomized trees (FS3), univariate FS using ANOVA (FS4) and RF (FS5), in addition to the intersection of markers selected by at least two of all FS methods (Inter1), the intersection of at least three of the three best FS methods (Inter2) and the intersection of the three best FS methods (Inter3).

| Model | All markers | FS1 | FS2 | FS3 | FS4 | FS5 | Inter1 | Inter2 | Inter3 |
|---|---|---|---|---|---|---|---|---|---|
| **AB** | 44.35 | 59.22 | 53.55 | 50.33 | 49.37 | 49.55 | 53.94 | 57.44 | 63.78 |
| **DT** | 39.22 | 59.02 | 48.39 | 48.07 | 44.83 | 45.97 | 50.93 | 55.72 | 61.02 |
| **GNB** | 44.9 | 64.82 | 85.67 | 58.19 | 62.29 | 53.39 | 76.88 | 86.17 | 69.91 |
| **GP** | 47.78 | 51.06 | 50.64 | 47.78 | 47.78 | 47.78 | 47.78 | 51.06 | 75.79 |
| **KNN** | 40.99 | 67.29 | 81.28 | 53.86 | 65.57 | 55.45 | 76.09 | 86.43 | 71.67 |
| **MLP** | 41.43 | 69.18 | 97.64 | 65.38 | 84.89 | 60.05 | 94.57 | 95.68 | 78.72 |
| **RF** | 48 | 76.1 | 79.58 | 65 | 64.63 | 61.89 | 74.98 | 79.52 | 71.71 |
| **SVM** | 49.58 | 72.35 | 88.59 | 63.01 | 77.72 | 59.39 | 84.28 | 87.82 | 73.48 |

**Supplementary Table 12. Cross-validation results of the predictive accuracy of each machine learning (ML) model and feature selection (FS) combination employed to predict clustering by SCYLV titer determined by RT-qPCR and SCYL symptom severity (SQ).** The machine learning models tested were adaptive boosting (AB), decision tree (DT), Gaussian naive Bayes (GNB), Gaussian process (GP), K-nearest neighbor (KNN), multilayer perceptron (MLP), random forest (RF) and support vector machine (SVM). FS methods tested were gradient tree boosting (FS1), L1-based FS through a linear support vector classification system (FS2), extremely randomized trees (FS3), univariate FS using ANOVA (FS4) and RF (FS5), in addition to the intersection of markers selected by at least two of all FS methods (Inter1), the intersection of at least three of the three best FS methods (Inter2) and the intersection of the three best FS methods (Inter3).

| Model | All markers | FS1 | FS2 | FS3 | FS4 | FS5 | Inter 1 | Inter 2 | Inter 3 |
|---|---|---|---|---|---|---|---|---|---|
| **AB** | 69.82 | 75.05 | 74.91 | 72.3 | 71.96 | 70.86 | 72.55 | 74.24 | 75.39 |
| **DT** | 64.92 | 80.48 | 73.1 | 70.03 | 69.92 | 64.72 | 73.42 | 75.71 | 83.08 |
| **GNB** | 73.81 | 78.84 | 82.2 | 70.34 | 79.13 | 67.71 | 82.75 | 87.66 | 69.32 |
| **GP** | 7.91 | 10.82 | 11.73 | 7.91 | 7.91 | 7.91 | 7.91 | 11.73 | 78.68 |
| **KNN** | 64.45 | 69.32 | 75.66 | 72.95 | 74.1 | 72.36 | 75.28 | 77.67 | 78.2 |
| **MLP** | 36.32 | 71.07 | 96.48 | 66.51 | 88.19 | 60.94 | 91.19 | 95.42 | 84.19 |
| **RF** | 73.13 | 78.12 | 74.8 | 73.93 | 74.5 | 73.91 | 76.26 | 77.42 | 85.4 |
| **SVM** | 73.92 | 73.92 | 76.51 | 73.92 | 73.01 | 73.92 | 75.51 | 86.49 | 84.55 |

**Supplementary Table 13. Markers selected by at least two of the three best feature selection methods, considering clustering by SCYLV titer determined by RT-qPCR (Q).** For each marker, the polymorphism (reference/alternative alleles) is provided.

| Marker | Polymorphism | Marker | Polymorphism |
|---|---|---|---|
| ECO_ACA_700_2-MSP_ACC-0201 | 0/1 | S4_5037630 | C/A |
| ECO_ACG_700_3-MSP_ACA-0350 | 0/1 | S4_51172780 | C/T |
| ECO_ACG_700_3-MSP_CGC-0192 | 0/1 | S4_55959846 | G/A |
| MSP_ACC-ECO_ACC-700-0487 | 0/1 | S4_62067807 | G/C |
| MSP_ACT-ECO_ACC-700-0165 | 0/1 | S4_62094357 | A/- |
| MSP_ACT-ECO_ACC-700-0185 | 0/1 | S4_72511101 | G/C |
| MSP_ACT-ECO_AGG-800-0116 | 0/1 | S5_16631107 | A/- |
| MSP_TAG-ECO_ACG_700_3-0186 | 0/1 | S5_19865296 | A/T |
| MSP_TTG-ECO_ACA-700-0133 | 0/1 | S5_19865299 | A/- |
| MSP_TTG-ECO_ACA-700-0135 | 0/1 | S5_30396509 | A/G |
| MSP_TTG-ECO_ACA-700-0238 | 0/1 | S5_37031999 | C/- |
| SCB381.09 | 0/1 | S5_37032002 | C/- |
| S1_10441059 | A/- | S5_37032003 | C/- |
| S1_1129066 | G/A | S5_37032005 | G/- |
| S1_13512336 | T/C | S5_37032006 | C/- |
| S1_3175304 | T/C | S5_37032007 | G/- |
| S1_33374268 | T/G | S5_37929370 | A/G |
| S1_33429745 | T/C | S5_38045669 | C/T |
| S1_33714180 | A/- | S5_39262786 | G/C |
| S1_33714200 | A/- | S5_39778394 | G/C |
| S1_36913604 | C/T | S5_4018386 | G/C |
| S1_38610862 | A/G | S5_40247302 | C/G |
| S1_39200477 | A/C | S5_51822173 | T/C |
| S1_3951700 | C/T | S5_58262048 | C/T |
| S1_50968980 | T/- | S5_58346648 | C/T |
| S1_73987041 | C/- | S5_64114690 | A/- |
| S1_75517043 | T/- | S5_70115114 | A/G |
| S1_78580392 | G/A | S5_77911159 | C/G |
| S1_80885415 | -/A | S5_79730136 | A/- |
| S1_84276876 | -/T | S5_81637789 | T/A |
| S1_85040557 | T/C | S5_83270565 | T/C |
| S1_89312901 | T/A | S5_86641489 | G/C |
| S1_89317063 | C/- | S5_87002893 | G/T |
| S1_90526407 | A/G | S5_87002894 | A/G |
| S1_94787735 | T/C | S5_87002896 | C/A |
| S2_101252947 | -/A | S5_87002901 | A/T |
| S2_101645661 | T/A | S6_23313127 | T/C |

| | | | |
|---|---|---|---|
| S2_102692127 | -/C | S6_31110903 | A/G |
| S2_10357059 | T/- | S6_37262886 | G/T |
| S2_103759601 | C/A | S6_39996328 | A/- |
| S2_104765621 | A/C | S6_45993856 | T/- |
| S2_107167277 | C/T | S6_47989239 | G/A |
| S2_107531177 | C/T | S6_49935949 | T/C |
| S2_11407928 | T/- | S6_50045923 | G/C |
| S2_11407929 | C/- | S6_51715201 | A/G |
| S2_114931317 | T/- | S6_54218288 | C/T |
| S2_117480270 | A/G | S6_7789087 | C/- |
| S2_11992364 | C/T | S6_7789088 | C/- |
| S2_22694123 | C/T | S6_88185776 | T/- |
| S2_31743348 | T/A | S6_90911017 | G/A |
| S2_3441185 | C/G | S6_94271628 | A/C |
| S2_4041569 | -/C | S6_97767289 | -/C |
| S2_47286302 | C/A | S7_16860739 | C/- |
| S2_580272 | T/C | S7_24246331 | C/T |
| S2_67409332 | C/- | S7_25494326 | T/A |
| S2_67409335 | T/- | S7_28163260 | G/C |
| S2_67409336 | T/- | S7_33839422 | G/A |
| S2_67409337 | C/- | S7_41636976 | A/G |
| S2_74145864 | G/C | S7_49705297 | G/T |
| S2_79667911 | A/G | S7_51204011 | A/- |
| S2_81096972 | A/T | S7_51204013 | C/- |
| S2_84707411 | A/G | S7_52954378 | G/A |
| S2_84898011 | G/C | S7_61409653 | C/T |
| S2_86412851 | C/G | S7_76845127 | C/T |
| S2_95211978 | A/- | S7_8727482 | A/C |
| S2_99027215 | T/C | S8_15175965 | T/- |
| S3_12466029 | T/A | S8_17563520 | T/G |
| S3_31119563 | -/T | S8_21423807 | A/C |
| S3_35454812 | -/G | S8_22051935 | -/T |
| S3_36459728 | -/G | S8_32042389 | A/G |
| S3_38170436 | C/A | S8_35612220 | G/- |
| S3_41060075 | T/A | S8_3868588 | C/A |
| S3_43081503 | A/T | S8_43975929 | G/A |
| S3_49105147 | G/A | S8_49191873 | C/T |
| S3_50218210 | T/- | S8_62846201 | G/A |
| S3_5029059 | A/- | S8_65429134 | A/- |
| S3_55613831 | T/- | SS_108682254 | A/T |
| S3_57815128 | C/A | SS_115845288 | -/A |
| S3_59564236 | A/- | SS_125997282 | G/C |
| S3_69969353 | C/- | SS_162306126 | A/G |

| | | | |
|---|---|---|---|
| S3_72076395 | C/T | SS_183828723 | A/G |
| S3_73057726 | -/C | SS_196174663 | C/T |
| S3_73057731 | C/- | SS_213100017 | A/G |
| S3_77376342 | T/A | SS_214099396 | G/- |
| S3_77376349 | C/G | SS_214099400 | -/T |
| S3_9810328 | A/G | SS_229367801 | A/G |
| S4_16437374 | T/C | SS_4267291 | T/G |
| S4_16582952 | A/G | SS_53385994 | G/C |
| S4_17019688 | T/C | SS_53417038 | G/A |
| S4_21976556 | C/T | SS_53615857 | T/- |
| S4_22573652 | A/G | SS_60091719 | G/- |
| S4_29702262 | C/T | SS_60091744 | -/C |
| S4_33241614 | -/G | SS_6997624 | A/- |
| S4_3664938 | A/T | SS_6997635 | -/A |
| S4_4503306 | C/T | SS_6997636 | -/A |

**Supplementary Table 14. Markers selected by at least two of the three best feature selection methods, considering clustering by SCYLV titer determined by RT-qPCR and SCYL symptom severity (SQ).** For each marker, the polymorphism (reference/alternative alleles) is provided.

| Marker | Polymorphism | Marker | Polymorphism |
|---|---|---|---|
| MSP_ACA-ECO_AAG_800_2-0121 | 0/1 | S3_77376349 | C/G |
| MSP_ACT-ECO_ACC-700-0104 | 0/1 | S4_15496266 | A/C |
| MSP_ACT-ECO_AGG-800-0106 | 0/1 | S4_17037843 | T/G |
| MSP_TTG-ECO_ACA-700-0094 | 0/1 | S4_20005282 | C/G |
| MSP_TTG-ECO_ACA-700-0133 | 0/1 | S4_21970615 | G/T |
| MSP_TTG-ECO_ACA-700-0238 | 0/1 | S4_21970622 | A/- |
| S1_106155638 | T/- | S4_26164754 | -/G |
| S1_106155646 | G/T | S4_27872858 | G/- |
| S1_106155651 | G/T | S4_29702516 | G/- |
| S1_1129066 | G/A | S4_38998631 | C/T |
| S1_13277038 | A/T | S4_52361599 | C/T |
| S1_15193842 | A/G | S4_60441000 | G/T |
| S1_15204569 | A/G | S4_67834354 | C/T |
| S1_16467489 | -/T | S4_71472623 | T/C |
| S1_21768021 | G/A | S4_72511101 | G/C |
| S1_27017708 | G/A | S5_12511335 | -/C |
| S1_32004666 | G/A | S5_34032519 | G/- |
| S1_33436946 | C/- | S5_34032520 | A/- |
| S1_33436948 | C/- | S5_41180286 | A/- |
| S1_36984611 | A/C | S5_62628785 | C/G |
| S1_55352018 | C/T | S5_64114690 | A/- |
| S1_6432921 | C/T | S5_70114127 | T/- |
| S1_6432925 | -/A | S5_70695514 | A/C |
| S1_6432926 | A/- | S5_74260966 | G/- |
| S1_73330330 | A/G | S5_78501954 | -/A |
| S1_73898167 | A/- | S5_83270565 | T/C |
| S1_75976908 | C/T | S5_85563234 | A/G |
| S1_76405126 | C/- | S6_16520022 | -/T |
| S1_81690247 | C/- | S6_21026466 | A/- |
| S1_96887191 | -/C | S6_21026467 | G/- |
| S1_98588359 | A/G | S6_21979304 | G/A |
| S2_102692127 | -/C | S6_43117749 | C/T |
| S2_10292064 | T/A | S6_44425039 | C/A |
| S2_10491272 | -/T | S6_51747509 | G/T |
| S2_107167277 | C/T | S6_69184546 | A/G |
| S2_10984287 | -/G | S7_14264782 | A/C |
| S2_12543228 | G/A | S7_21392489 | C/T |

| | | | |
|---|---|---|---|
| S2_1695387 | G/A | S7_25558360 | T/G |
| S2_20752623 | -/G | S7_37524792 | T/C |
| S2_2344737 | A/G | S7_40187897 | T/C |
| S2_29082911 | T/C | S7_4925034 | T/C |
| S2_32500210 | -/T | S7_7265178 | T/C |
| S2_3384263 | -/C | S7_78538330 | G/A |
| S2_36266547 | -/T | S8_15352646 | A/C |
| S2_62995081 | G/A | S8_15352661 | C/T |
| S2_67466872 | G/A | S8_17563520 | T/G |
| S2_71898215 | C/T | S8_20515788 | A/C |
| S2_78043402 | C/A | S8_26548878 | C/G |
| S2_80240920 | A/T | S8_39882618 | G/T |
| S2_85812601 | T/- | S8_41932797 | A/C |
| S3_19279325 | G/C | S8_50905917 | C/T |
| S3_2171611 | -/A | S8_59270955 | A/G |
| S3_43081503 | A/T | S8_8575896 | T/G |
| S3_49665310 | A/G | SS_100090050 | A/T |
| S3_5029059 | A/- | SS_185211457 | A/G |
| S3_53391124 | G/C | SS_190207740 | A/G |
| S3_53603607 | A/C | SS_28567348 | G/A |
| S3_54707120 | T/G | SS_53615857 | T/- |
| S3_55712812 | T/C | SS_53991168 | C/T |
| S3_74204066 | G/- | SS_84033755 | A/G |

**Supplementary Table 15. Annotation of markers associated with the three analyzed traits.** Annotation was obtained by aligning 2,000-bp neighboring regions with the genomes of closely related plant species. For each marker, the approach used for association, associated trait, gene, alignment percentage of identity (% ID), alignment E-value and gene ontology annotation are provided.

| Marker | Approach | Gene | %ID | E-value | Annotation |
|---|---|---|---|---|---|
| SCB381.09 | ML_SQ | SbRio.10G317500.1 | 95.55 | 0 | Peroxidase precursor, putative, expressed |
| S1_10441059 | ML_Q | Sobic.001G494500.2 | 97.61 | 5.51E-118 | Kinase, pfkB family, putative, expressed |
| S1_105811826 | GWAS_Rel | Sobic.001G053100.2 | 78.73 | 4.08E-149 | Expressed protein |
| S1_13277038 | ML_SQ | Oropetium_20150105_00333A | 97.17 | 2.20E-42 | Prenyltransferase, putative, expressed |
| S1_13512336 | ML_Q | SbRio.01G504900.1 | 99.07 | 7.86E-47 | OsSCP11 - Putative Serine Carboxypeptidase homologue, expressed |
| S1_15193842 | ML_SQ | Sobic.001G464100.1 | 98.83 | 3.48E-80 | Cytochrome b5-like Heme/Steroid binding domain containing protein, expressed |
| S1_15204569 | ML_SQ | Zm00001d048138_T026 | 97.85 | 7.98E-37 | BTBN4 - Bric-a-Brac, Tramtrack, Broad Complex BTB domain with nonphototropic hypocotyl 3 NPH3 domain, expressed |
| S1_16467489 | ML_SQ | Sobic.001G452600.1 | 98.47 | 7.02E-127 | Leucine-rich repeat family protein, putative, expressed |
| S1_19840994 | GWAS_Sym | Sobic.001G417400.1 | 94.54 | 5.66E-98 | RNA recognition motif containing protein, putative, expressed |
| S1_3175304 | ML_Q | Sobic.001G532200.1 | 96.18 | 4.20E-129 | OsRhmbd7 - Putative Rhomboid homologue, expressed |
| S1_32004666 | ML_SQ | Sobic.001G346600.1 | 98.03 | 5.91E-68 | Kinesin heavy chain isolog, putative, expressed |
| S1_33374268 | ML_Q | Pavir.3KG265854.1 | 83.33 | 7.64E-67 | Transposon protein, putative, unclassified, expressed |
| S1_33436946 | ML_SQ | SbRio.01G362100.1 | 91.46 | 0 | Cytochrome P450, putative, expressed |
| S1_33436948 | ML_SQ | SbRio.01G362100.1 | 91.46 | 0 | Cytochrome P450, putative, expressed |
| S1_38610862 | ML_Q | Sobic.001G305100.1 | 98.23 | 6.08E-48 | Pectinacetylesterase domain containing protein, expressed |
| S1_3951700 | ML_Q | Sobic.001G528000.1 | 95.19 | 6.17E-38 | SNARE domain containing protein, putative, expressed |
| S1_55352018 | ML_SQ | SbRio.06G054100.1 | 99.03 | 4.73E-44 | Vacuolar-sorting receptor precursor, putative, expressed |
| S1_6432921 | ML_SQ | Zm00001d027607_T001 | 98.31 | 3.63E-50 | Cysteine-rich repeat secretory protein precursor, putative, expressed |

| | | | | | |
|---|---|---|---|---|---|
| S1_6432925 | ML_SQ | Zm00001d027607_T001 | 98.31 | 3.63E-50 | Cysteine-rich repeat secretory protein precursor, putative, expressed |
| S1_6432926 | ML_SQ | Zm00001d027607_T001 | 98.31 | 3.63E-50 | Cysteine-rich repeat secretory protein precursor, putative, expressed |
| S1_7029466 | GWAS_Sym | SbRio.01G540100.1 | 94.51 | 0 | GATA zinc finger domain containing protein, expressed |
| S1_7029472 | GWAS_Sym | SbRio.01G540100.1 | 94.61 | 0 | GATA zinc finger domain containing protein, expressed |
| S1_7029473 | GWAS_Sym | SbRio.01G540100.1 | 94.61 | 0 | GATA zinc finger domain containing protein, expressed |
| S1_7029474 | GWAS_Sym | SbRio.01G540100.1 | 94.61 | 0 | GATA zinc finger domain containing protein, expressed |
| S1_7029475 | GWAS_Sym | SbRio.01G540100.1 | 94.61 | 0 | GATA zinc finger domain containing protein, expressed |
| S1_7029479 | GWAS_Sym | SbRio.01G540100.1 | 94.58 | 0 | GATA zinc finger domain containing protein, expressed |
| S1_7029480 | GWAS_Sym | SbRio.01G540100.1 | 94.57 | 0 | GATA zinc finger domain containing protein, expressed |
| S1_7029481 | GWAS_Sym | SbRio.01G540100.1 | 94.56 | 0 | GATA zinc finger domain containing protein, expressed |
| S1_7029482 | GWAS_Sym | SbRio.01G540100.1 | 94.55 | 0 | GATA zinc finger domain containing protein, expressed |
| S1_7029483 | GWAS_Sym | SbRio.01G540100.1 | 94.54 | 0 | GATA zinc finger domain containing protein, expressed |
| S1_73330330 | ML_SQ | Sobic.001G217200.2 | 98.94 | 4.44E-89 | START domain containing protein, expressed |
| S1_73898167 | ML_SQ | Sobic.001G214000.1 | 98.48 | 5.17E-163 | Dicer, putative, expressed |
| S1_75976908 | ML_SQ | SbRio.01G476900.1 | 100.00 | 2.87E-36 | AP2/EREBP transcription factor BABY BOOM, putative, expressed |
| S1_76405126 | ML_SQ | Zm00001d032575_T013 | 97.70 | 1.73E-33 | Expressed protein |
| S1_77494328 | GWAS_Sym | Sobic.001G200200.2 | 97.80 | 1.03E-35 | Tetratricopeptide repeat, putative, expressed |
| S1_77494331 | GWAS_Sym | Sobic.001G200200.2 | 97.80 | 1.03E-35 | Tetratricopeptide repeat, putative, expressed |
| S1_78580392 | ML_Q | Sobic.001G192000.1 | 96.32 | 0 | Autophagy protein 9, putative, expressed |
| S1_80885415 | ML_Q | Sobic.001G186700.1 | 94.25 | 0 | Zinc finger C-x8-C-x5-C-x3-H type family protein |
| S1_81690247 | ML_SQ | Sobic.001G185200.1 | 96.43 | 4.73E-44 | Expressed protein |
| S1_85040557 | ML_Q | Sobic.001G163300.2 | 96.23 | 0 | Zinc finger protein-related, putative, expressed |
| S1_90526407 | ML_Q | Sobic.001G122900.1 | 95.68 | 1.00E-55 | Expressed protein |
| S1_96887191 | ML_SQ | Sobic.001G106500.1 | 88.68 | 5.95E-63 | Expressed protein |

| | | | | | |
|---|---|---|---|---|---|
| S2_100207393 | GWAS_Rel | Sobic.008G107100.1 | 93.23 | 0 | Cytochrome P450, putative, expressed |
| S2_101645661 | ML_Q | SbRio.08G127900.1 | 95.20 | 6.08E-48 | rhoGAP domain containing protein, expressed |
| S2_102692127 | ML_Q/ML_SQ | Seita.6G074900.1 | 91.20 | 7.81E-52 | BTBN17 - Bric-a-Brac, Tramtrack, Broad Complex BTB domain with nonphototropic hypocotyl 3 NPH3 and coiled-coil domains, expressed |
| S2_10357059 | ML_Q | GRMZM2G017080_T02 | 96.15 | 4.77E-39 | Hydrolase protein, putative, expressed |
| S2_104765621 | ML_Q | Sobic.008G132550.1 | 87.97 | 0 | Pleiotropic drug resistance protein, putative, expressed |
| S2_10491272 | ML_SQ | Sobic.002G380700.1 | 94.82 | 1.19E-114 | Expressed protein |
| S2_107167277 | ML_Q/ML_SQ | Sobic.008G141100.1 | 100.00 | 4.73E-44 | Retrotransposon protein, putative, unclassified, expressed |
| S2_111395374 | GWAS_Rel | Sobic.008G156600.1 | 95.53 | 9.35E-106 | Leucine Rich Repeat family protein, expressed |
| S2_12543228 | ML_SQ | Sobic.002G363500.2 | 100.00 | 4.77E-39 | RNA methyltransferase, TrmH family protein, putative, expressed |
| S2_1695387 | ML_SQ | Zm00001d007241_T001 | 97.94 | 4.77E-39 | KIP1, putative, expressed |
| S2_20752623 | ML_SQ | Sobic.002G312800.1 | 97.75 | 9.68E-81 | WD domain, G-beta repeat domain containing protein, expressed |
| S2_2344737 | ML_SQ | Sobic.002G426800.1 | 93.87 | 0 | Zinc finger, C3HC4 type domain containing protein, expressed |
| S2_29082911 | ML_SQ | Sobic.002G259000.1 | 99.02 | 5.66E-98 | Ribosomal protein L7Ae, putative, expressed |
| S2_31743348 | ML_Q | SbRio.02G255400.1 | 93.50 | 0 | Nucleolar GTP-binding protein 1, putative, expressed |
| S2_32500210 | ML_SQ | Sobic.002G237800.2 | 100.00 | 3.68E-40 | NOL1/NOP2/sun family protein, putative, expressed |
| S2_36266547 | ML_SQ | Pavir.2NG346303.1 | 94.18 | 4.29E-114 | Expressed protein |
| S2_4041569 | ML_Q | Sobic.002G415100.2 | 92.18 | 5.95E-63 | POEI50 - Pollen Ole e I allergen and extensin family protein precursor, expressed |
| S2_580272 | ML_Q | Thint.06G0495000.1 | 96.77 | 1.01E-50 | MYB family transcription factor, putative, expressed |
| S2_62995081 | ML_SQ | Sobic.002G124900.1 | 97.12 | 2.85E-41 | ATP synthase D chain, mitochondrial |
| S2_67409332 | ML_Q | Sobic.002G108000.1 | 94.22 | 7.38E-92 | Myosin, putative, expressed |
| S2_67409337 | ML_Q | Sobic.002G108000.1 | 94.22 | 7.38E-92 | Myosin, putative, expressed |
| S2_67466872 | ML_SQ | Sobic.002G106700.1 | 91.85 | 0 | OsWAK75 - OsWAK receptor-like protein kinase, expressed |
| S2_74145864 | ML_Q | Sobic.002G081500.5 | 92.17 | 1.67E-58 | Glycerol-3-phosphate dehydrogenase, putative, expressed |

| | | | | | |
|---|---|---|---|---|---|
| S2_78043402 | ML_SQ | SbRio.02G066000.1 | 97.22 | 1.57E-98 | OsSub3 - Putative Subtilisin homologue, expressed |
| S2_79667911 | ML_Q | SbRio.02G057300.1 | 89.06 | 0 | Rf1, mitochondrial precursor, putative, expressed |
| S2_85812601 | ML_SQ | Sobic.002G019750.1 | 88.94 | 2.11E-72 | OsFBX234 - F-box domain containing protein, expressed |
| S2_86412851 | ML_Q | SbRio.02G018600.1 | 92.54 | 3.31E-115 | Expressed protein |
| S3_19279325 | ML_SQ | Sobic.003G287000.1 | 99.09 | 6.08E-48 | BURP domain containing protein, expressed |
| S3_2171611 | ML_SQ | Sobic.003G395400.1 | 99.30 | 1.48E-143 | BTBZ2 - Bric-a-Brac, Tramtrack, and Broad Complex BTB domain with TAZ zinc finger and Calmodulin-binding domains, expressed |
| S3_35454812 | ML_Q | Zm00001d011212_T001 | 93.14 | 1.73E-33 | Helix-loop-helix DNA-binding domain containing protein, expressed |
| S3_41060075 | ML_Q | Brasy7G095300.1 | 89.72 | 0 | Expressed protein |
| S3_48163530 | GWAS_Sym | SbRio.03G158900.1 | 86.38 | 0 | OsFBX103 - F-box domain containing protein, expressed |
| S3_49105147 | ML_Q | Sobic.003G146900.1 | 99.20 | 2.79E-56 | Lipase class 3 family protein, putative, expressed |
| S3_50218210 | ML_Q | SbRio.03G148100.1 | 97.24 | 0 | Receptor-like protein kinase 5 precursor, putative, expressed |
| S3_54707120 | ML_SQ | SbRio.03G120100.1 | 97.14 | 0 | Lipase, putative, expressed |
| S3_54754895 | GWAS_Sym | SbRio.03G120000.1 | 93.77 | 5.13E-168 | GCRP6 - Glycine and cysteine rich family protein precursor, putative, expressed |
| S3_55613831 | ML_Q | Zm00001d040210_T001 | 98.12 | 3.38E-100 | Protein of unknown function (DUF 3339) |
| S3_57815128 | ML_Q | Sobic.003G101500.1 | 93.75 | 0 | DNAJ domain containing protein, expressed |
| S3_72076395 | ML_Q | Oropetium_20150105_17951A | 97.39 | 2.19E-47 | BES1/BZR1 homolog protein, putative, expressed |
| S3_74204066 | ML_SQ | Sobic.003G013100.1 | 92.57 | 2.17E-52 | GDSL-like lipase/acylhydrolase, putative, expressed |
| S4_11629587 | GWAS_Sym | Sobic.004G242500.1 | 95.95 | 9.96E-61 | Glycosyl hydrolase family 47 domain contain protein, expressed |
| S4_15496266 | ML_SQ | Sobic.004G267400.2 | 98.59 | 4.60E-64 | Basic helix-loop-helix, putative, expressed |
| S4_16437374 | ML_Q | Sobic.004G274000.1 | 95.94 | 0 | STE_MEK_ste7_MAP2K.4 - STE kinases include homologs to sterile 7, sterile 11 and sterile 20 from yeast, expressed |
| S4_16582952 | ML_Q | Zm00001d017699_T001 | 92.08 | 1.63E-73 | Glycosyl hydrolases family 16, putative, expressed |
| S4_17037843 | ML_SQ | Sobic.004G276900.2 | 98.14 | 1.87E-157 | Leucyl-tRNA synthetase, cytoplasmic, putative, expressed |
| S4_20005282 | ML_SQ | Sobic.004G296500.2 | 95.58 | 6.12E-43 | Serine esterase family protein, putative, expressed |
| S4_21970615 | ML_SQ | Sobic.004G224350.1 | 90.41 | 2.62E-101 | Stage II sporulation protein E, putative, expressed |

| | | | | | |
|---|---|---|---|---|---|
| S4_21970622 | ML_SQ | Sobic.004G224350.1 | 90.64 | 3.36E-105 | Stage II sporulation protein E, putative, expressed |
| S4_26164754 | ML_SQ | Pahal.1G292400.1 | 97.30 | 3.66E-45 | Expressed protein |
| S4_29702262 | ML_Q | SbRio.04G200400.1 | 98.14 | 1.17E-129 | Transporter family protein, putative, expressed |
| S4_29702516 | ML_SQ | SbRio.04G200400.1 | 98.14 | 1.17E-129 | Transporter family protein, putative, expressed |
| S4_33241614 | ML_Q | Sobic.004G165600.1 | 92.04 | 1.33E-34 | Glycosyl hydrolases family 17, putative, expressed |
| S4_3664938 | ML_Q | SbRio.04G366500.1 | 96.36 | 9.28E-111 | Uncharacterized protein yqjG, putative, expressed |
| S4_38998631 | ML_SQ | Sobic.004G148100.1 | 95.76 | 1.02E-45 | Cupin domain containing protein, expressed |
| S4_4503306 | ML_Q | Zm00001d018419_T001 | 95.76 | 1.02E-45 | Expressed protein |
| S4_5037630 | ML_Q | Sobic.004G333100.1 | 98.85 | 7.48E-82 | Expressed protein |
| S4_52361599 | ML_SQ | Sobic.003G167000.1 | 89.81 | 4.70E-49 | Protein phosphotase protein, putative, expressed |
| S4_60441000 | ML_SQ | Sobic.004G070800.1 | 98.86 | 4.20E-129 | TIC21, putative, expressed |
| S4_62067807 | ML_Q | Zm00001d053733_T001 | 80.98 | 2.07E-87 | Glycosyl hydrolases family 17, putative, expressed |
| S4_62622513 | GWAS_Sym | HORVU6Hr1G031450.3 | 97.09 | 1.02E-40 | OsSPL4 - SBP-box gene family member, expressed |
| S4_67834354 | ML_SQ | SbRio.01G107700.1 | 94.32 | 0 | gp176, putative, expressed |
| S4_71472623 | ML_SQ | Sobic.004G014100.1 | 98.35 | 1.24E-84 | Sialyltransferase family domain containing protein, expressed |
| S4_72511101 | ML_Q/ML_SQ | HORVU2Hr1G108070.13 | 97.73 | 4.80E-34 | Glycosyltransferase protein, putative, expressed |
| S5_30396509 | ML_Q | Zm00001d025474_T003 | 98.37 | 5.51E-118 | Chloride channel C |
| S5_34032519 | ML_SQ | Sobic.006G074900.1 | 95.03 | 0 | Transposon protein, putative, Mutator subclass, expressed |
| S5_34032520 | ML_SQ | Sobic.006G074900.1 | 95.02 | 0 | Transposon protein, putative, Mutator subclass, expressed |
| S5_38045669 | ML_Q | Sobic.006G058800.1 | 99.49 | 0 | tRNA synthetase class II core domain containing protein, expressed |
| S5_51822173 | ML_Q | SbRio.06G023600.1 | 98.36 | 1.98E-117 | ABC transporter family protein, putative, expressed |
| S5_58262048 | ML_Q | SbRio.06G005000.1 | 94.00 | 2.02E-102 | Erythronate-4-phosphate dehydrogenase domain containing protein, expressed |
| S5_70695514 | ML_SQ | Zm00001d049007_T001 | 97.86 | 3.34E-110 | PHD finger protein, putative, expressed |
| S5_83270565 | ML_Q/ML_SQ | Sobic.005G039500.1 | 100.00 | 2.19E-47 | Signal recognition particle 54 kDa protein, putative, expressed |
| S5_85563234 | ML_SQ | Sobic.005G029100.2 | 98.79 | 7.54E-77 | PP2A regulatory subunit TAP46, putative, expressed |
| S5_87002893 | ML_Q | Sobic.005G020700.1 | 96.70 | 4.80E-34 | MATE efflux family protein, putative, expressed |

| | | | | | |
|---|---|---|---|---|---|
| S5_87002894 | ML_Q | Sobic.005G020700.1 | 96.70 | 4.80E-34 | MATE efflux family protein, putative, expressed |
| S5_87002896 | ML_Q | Sobic.005G020700.1 | 96.70 | 4.80E-34 | MATE efflux family protein, putative, expressed |
| S5_87002901 | ML_Q | Sobic.005G020700.1 | 96.70 | 4.80E-34 | MATE efflux family protein, putative, expressed |
| S6_16520022 | ML_SQ | Sobic.007G152000.1 | 98.77 | 8.03E-32 | Activator of 90 kDa heat shock protein ATPase homolog, putative, expressed |
| S6_19432542 | GWAS_Sym | Pahal.6G214300.1 | 81.90 | 7.92E-42 | STE_MEK_ste7_MAP2K.8 - STE kinases include homologs to sterile 7, sterile 11 and sterile 20 from yeast, expressed |
| S6_20347039 | GWAS_Sym | Pahal.6G208600.1 | 96.65 | 5.58E-108 | Copper transport protein family |
| S6_39996328 | ML_Q | Sobic.009G009500.1 | 92.89 | 7.43E-87 | Inhibitor I family protein, putative, expressed |
| S6_43117749 | ML_SQ | Sobic.007G092500.1 | 86.81 | 7.22E-107 | Expressed protein |
| S6_45993856 | ML_Q | Sobic.007G085400.1 | 90.85 | 0 | NBS-LRR disease resistance protein, putative, expressed |
| S6_47989239 | ML_Q | Sobic.007G079300.1 | 96.73 | 0 | Proteasome/cyclosome repeat containing protein, expressed |
| S6_49935949 | ML_Q | Sobic.007G073700.1 | 97.12 | 0 | Nucleoside transporter, putative, expressed |
| S6_50045923 | ML_Q | Sobic.007G073400.1 | 97.37 | 9.61E-86 | SHR5-receptor-like kinase, putative, expressed |
| S6_51715201 | ML_Q | Sobic.007G069700.1 | 98.96 | 3.68E-40 | Nucleolin, putative, expressed |
| S6_69184546 | ML_SQ | Sobic.007G007000.1 | 98.13 | 7.59E-72 | DUF250 domain containing protein, putative, expressed |
| S6_7789087 | ML_Q | Sobic.007G198400.1 | 87.65 | 6.97E-132 | BTB/POZ domain containing protein, expressed |
| S6_7789088 | ML_Q | Sobic.007G198400.1 | 87.65 | 6.97E-132 | BTB/POZ domain containing protein, expressed |
| S6_94271628 | ML_Q | Sobic.005G194000.2 | 96.21 | 6.04E-53 | DUF1399 containing protein, putative, expressed |
| S6_97767289 | ML_Q | Sobic.005G203500.1 | 87.07 | 0 | LZ-NBS-LRR class RGA, putative, expressed |
| S7_14264782 | ML_SQ | Sobic.009G174400.7 | 97.56 | 7.81E-52 | Protein of unknown function (DUF3133) |
| S7_16860739 | ML_Q | Sobic.009G161401.1 | 96.91 | 2.22E-37 | Expressed protein |
| S7_24246331 | ML_Q | Zm00001d038089_T002 | 98.11 | 4.73E-44 | Integral membrane transporter family protein, putative, expressed |
| S7_25132085 | GWAS_Abs | Sobic.009G121100.2 | 91.26 | 5.87E-73 | Plant calmodulin-binding protein-related |
| S7_25558360 | ML_SQ | Zm00001d037864_T030 | 97.58 | 1.63E-73 | Tetratricopeptide repeat containing protein, putative, expressed |
| S7_28163260 | ML_Q | Sobic.009G108700.2 | 96.81 | 1.03E-35 | NADP-dependent malic enzyme, chloroplast precursor, putative, expressed |
| S7_33839422 | ML_Q | GRMZM2G045892_T01 | 97.92 | 1.71E-38 | CBS domain containing protein, expressed |

| | | | | | |
|---|---|---|---|---|---|
| S7_40187897 | ML_SQ | Sobic.001G399200.2 | 89.36 | 7.92E-42 | Kinesin motor domain containing protein, expressed |
| S7_41636976 | ML_Q | Sobic.009G065200.2 | 96.52 | 0 | Zinc finger family protein, putative, expressed |
| S7_4925034 | ML_SQ | Sevir.3G147201.1 | 97.93 | 1.65E-63 | WRKY16, expressed |
| S7_49705297 | ML_Q | Sobic.009G035100.1 | 96.20 | 2.75E-66 | Thioesterase family protein, putative, expressed |
| S7_51204011 | ML_Q | GRMZM5G800780_T01 | 95.18 | 0 | Cytochrome b6, putative, expressed |
| S7_51204013 | ML_Q | GRMZM5G800780_T01 | 95.18 | 0 | Cytochrome b6, putative, expressed |
| S7_61409653 | ML_Q | Pahal.3G419900.1 | 88.98 | 1.33E-34 | UDP-glucoronosyl/UDP-glucosyl transferase, putative, expressed |
| S7_8727482 | ML_Q | Sobic.009G204800.1 | 97.86 | 2.77E-61 | Senescence-induced receptor-like serine/threonine-protein kinase precursor, putative, expressed |
| S8_10096401 | GWAS_Sym | Sobic.010G230300.1 | 96.75 | 3.63E-50 | Galactosyltransferase, putative, expressed |
| S8_10096402 | GWAS_Sym | Sobic.010G230300.1 | 96.77 | 1.01E-50 | Galactosyltransferase, putative, expressed |
| S8_10096403 | GWAS_Sym | Sobic.010G230300.1 | 96.80 | 2.81E-51 | Galactosyltransferase, putative, expressed |
| S8_10096406 | GWAS_Sym | Sobic.010G230300.1 | 96.88 | 6.04E-53 | Galactosyltransferase, putative, expressed |
| S8_10096428 | GWAS_Sym | Sobic.010G230300.1 | 97.33 | 3.56E-65 | Galactosyltransferase, putative, expressed |
| S8_15352646 | ML_SQ | Sobic.010G204400.1 | 99.23 | 4.63E-59 | Phox domain-containing protein, putative, expressed |
| S8_15352661 | ML_SQ | Sobic.010G204400.1 | 99.23 | 4.63E-59 | Phox domain-containing protein, putative, expressed |
| S8_17563520 | ML_Q/ML_SQ | Zm00001d046725_T001 | 92.48 | 5.62E-103 | D-mannose binding lectin family protein, expressed |
| S8_20515788 | ML_SQ | Sobic.010G180400.1 | 100.00 | 7.38E-92 | Myristoyl-acyl carrier protein thioesterase, chloroplast precursor, putative, expressed |
| S8_22051935 | ML_Q | Sobic.010G172600.1 | 93.20 | 0 | Expressed protein |
| S8_26548878 | ML_SQ | Sobic.010G160500.4 | 98.91 | 6.17E-38 | DEAD-box ATP-dependent RNA helicase, putative, expressed |
| S8_35612220 | ML_Q | Zm00001d052404_T002 | 87.50 | 3.71E-35 | ELMO/CED-12 family protein, putative, expressed |
| S8_39882618 | ML_SQ | Sobic.010G131300.2 | 97.79 | 1.17E-129 | ABB1 - Ankyrin repeat region with 2 Bric-a-Brac, Tramtrack, Broad Complex BTB domains, expressed |
| S8_41932797 | ML_SQ | Sobic.010G124300.1 | 96.77 | 2.04E-97 | Oxidoreductase, short chain dehydrogenase/reductase family domain containing protein, expressed |
| S8_59270955 | ML_SQ | SbRio.10G053600.1 | 98.21 | 7.54E-77 | Tyrosylprotein sulfotransferase |
| S8_62846201 | ML_Q | Sobic.010G031300.1 | 98.31 | 1.01E-50 | Auxin-induced protein 5NG4, putative, expressed |

| | | | | | |
|---|---|---|---|---|---|
| S8_8575896 | ML_SQ | Sobic.010G239600.2 | 98.45 | 2.79E-56 | RNA polymerase I specific transcription initiation factor RRN3 family protein, putative, expressed |
| SS_100090050 | ML_SQ | Zm00001d045565_T001 | 95.04 | 1.02E-45 | Phosphoglycolate phosphatase, plasmid, putative, expressed |
| SS_108682254 | ML_Q | Sobic.005G028100.1 | 99.17 | 4.67E-54 | Nucleolar complex protein 2, putative, expressed |
| SS_115845288 | ML_Q | Sobic.004G276700.1 | 92.48 | 1.32E-44 | Transposon protein, putative, unclassified, expressed |
| SS_185211457 | ML_SQ | Sobic.002G140800.2 | 98.75 | 2.74E-31 | RNA pseudouridine synthase, putative, expressed |
| SS_196174663 | ML_Q | Sobic.010G134500.1 | 97.60 | 6.04E-53 | Exonuclease, putative, expressed |
| SS_213100017 | ML_Q | SbRio.07G035500.1 | 91.32 | 0 | Pentatricopeptide repeat domain containing protein, putative, expressed |
| SS_229367801 | ML_Q | Sobic.002G296800.3 | 92.17 | 2.11E-37 | Cell cycle control protein, putative, expressed |
| SS_28567348 | ML_SQ | SbRio.05G003000.1 | 97.46 | 3.24E-130 | Double Clp-N motif-containing P-loop nucleoside triphosphate hydrolases superfamily protein |
| SS_4267291 | ML_Q | Sobic.004G290000.1 | 100.00 | 3.24E-95 | OsSub19 - Putative Subtilisin homologue, expressed |
| SS_53615857 | ML_Q/ML_SQ | Sobic.008G082000.1 | 92.31 | 1.31E-49 | RWP-RK domain-containing protein, putative, expressed |
| SS_84033755 | ML_SQ | Sevir.7G299100.4 | 84.07 | 2.61E-66 | Transposon protein, putative, Pong subclass, expressed |

**Supplementary Figures**

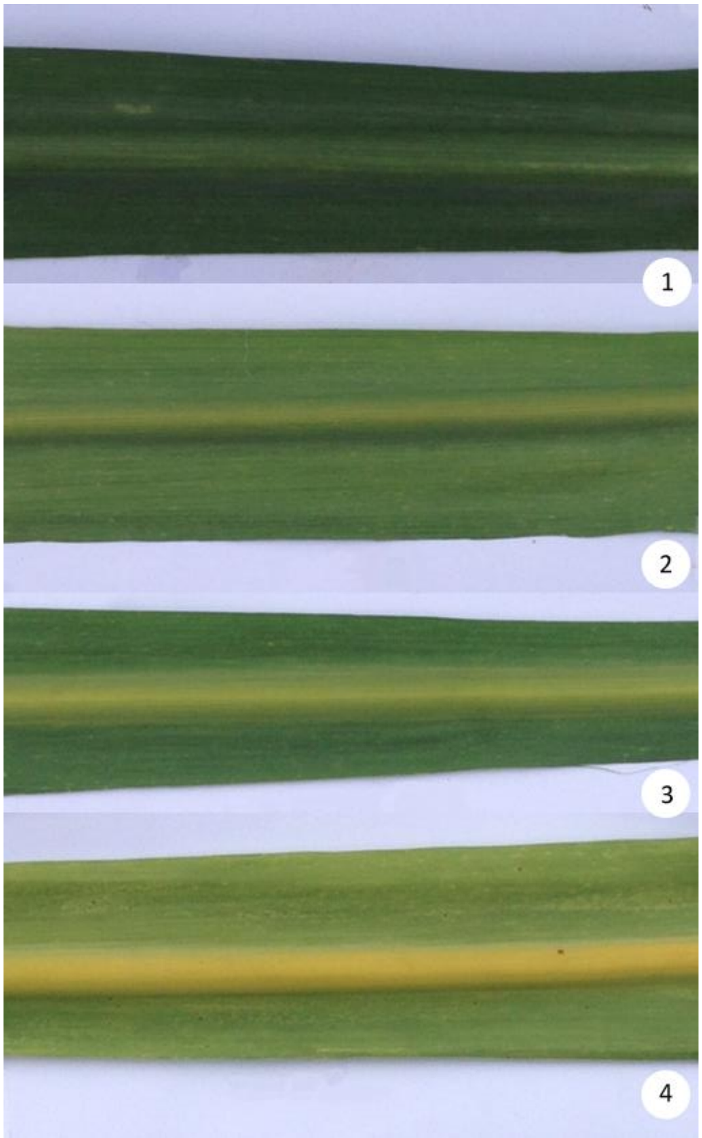


**Supplementary Fig. 1 Diagram of the scoring scale for the assessment of sugarcane yellow leaf symptom severity.** (1) Green leaf with no symptoms; (2) slight yellowing of the abaxial midrib, with a discrete or no progression to the leaf blade; (3) intense yellowing of the abaxial midrib and a more pronounced yellowing of the leaf blade; (4) intense yellowing of both abaxial midrib and leaf blade. Adapted from Burbano et al. (2021)

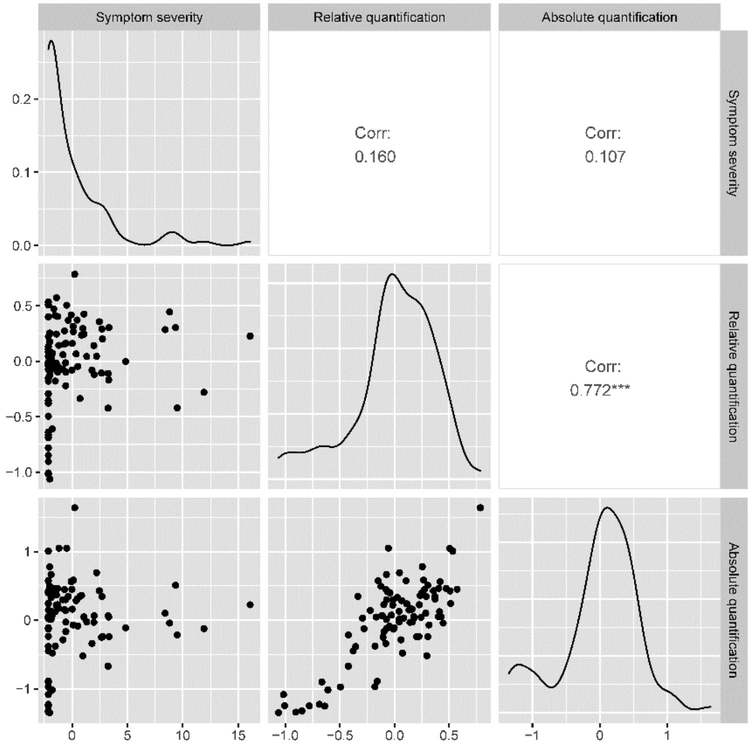


**Supplementary Fig. 2 Distributions of BLUP values and correlations between the three traits analyzed.**

Asterisks (***) indicate statistical significance ($p < 0.0005$)

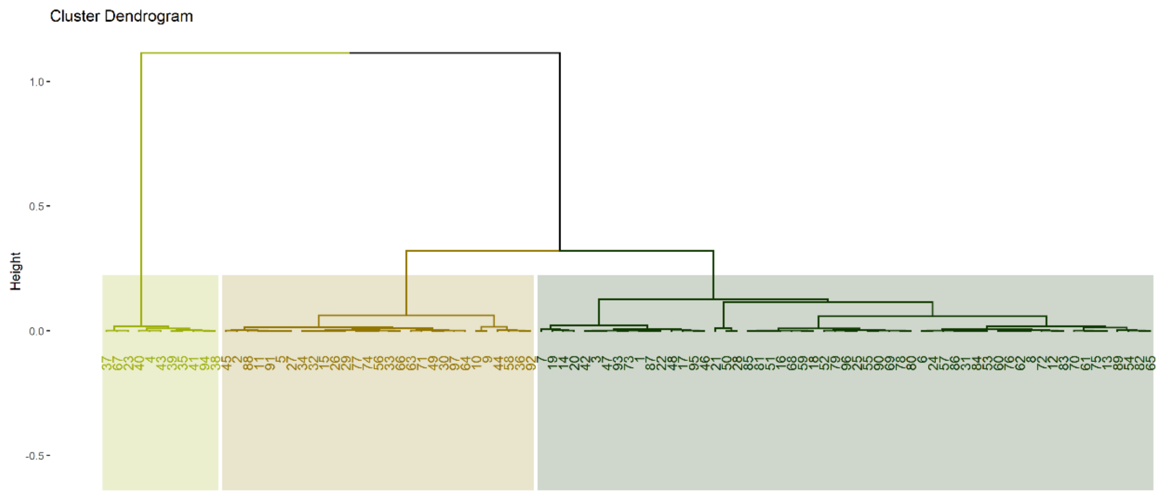


**Supplementary Fig. 3 Cluster dendrogram generated in the hierarchical clustering on principal components (HCPC) analysis using the BLUPs of SCYLV titer determined by RT-qPCR.** A division into three clusters (Q1/light green, Q2/brown and Q3/dark green) was considered

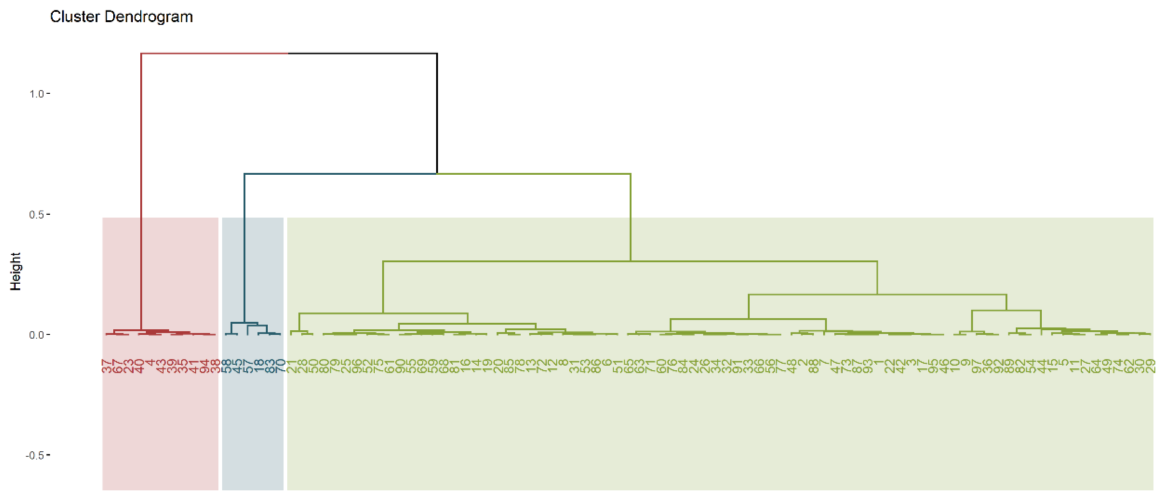


**Supplementary Fig. 4 Cluster dendrogram generated in the hierarchical clustering on principal components (HCPC) analysis using BLUPs of SCYL symptom severity and SCYLV titer determined by RT-qPCR.** A division into three clusters (SQ1/red, SQ2/green and SQ3/blue) was considered

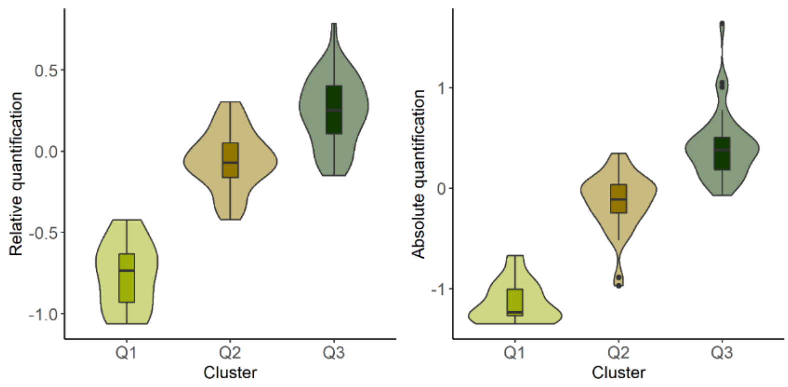


**Supplementary Fig. 5 Boxplots depicting the distributions of BLUP values of SCYLV titer determined by relative and absolute quantifications among the three clusters identified in the hierarchical clustering on principal components (HCPC) using the BLUPs of SCYLV titer determined by RT-qPCR.** A division into three clusters (Q1, Q2 and Q3) was considered

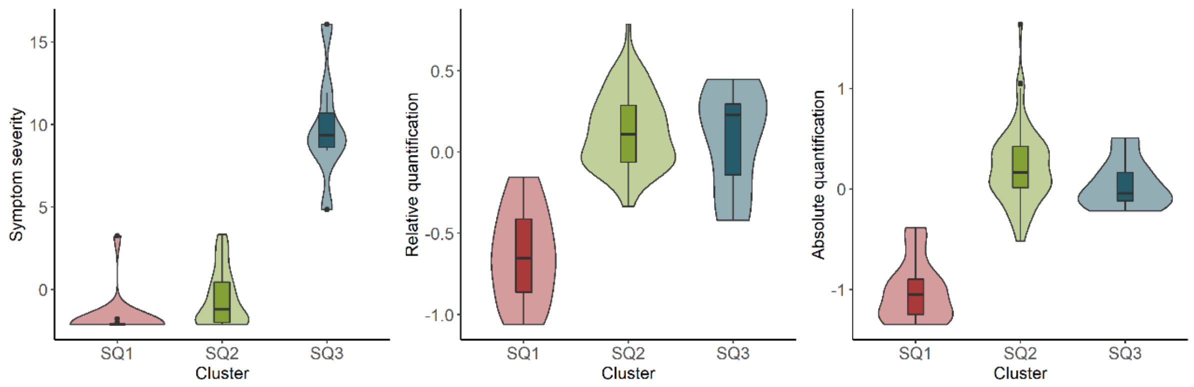


**Supplementary Fig. 6 Boxplots depicting the distributions of BLUP values for the three analyzed traits among the three clusters identified in the hierarchical clustering on principal components (HCPC) using BLUPs of SCYL symptom severity and SCYLV titer determined by RT-qPCR.** A division into three clusters (SQ1, SQ2 and SQ3) was considered

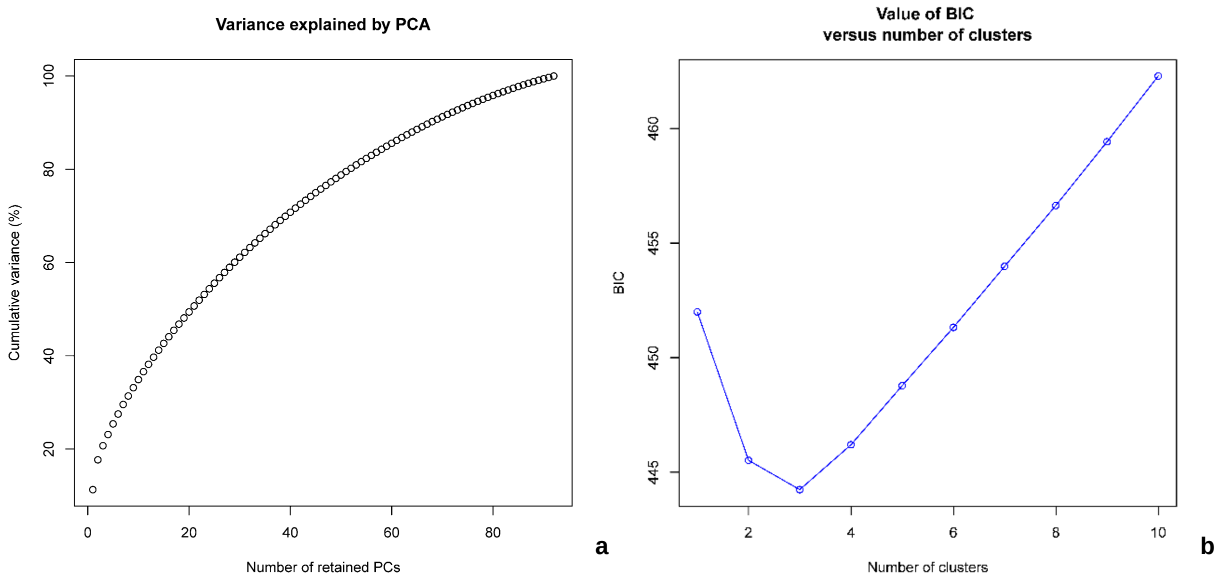


**Supplementary Fig. 7 Graphical output of methods employed for the determination of the optimal number of genetic clusters in the panel based on a discriminant analysis of principal components (DAPC) using 622 dominant markers. (a)** Cumulative variance explained by the principal components in the analysis. **(b)** Bayesian information criterion (BIC) versus number of clusters in k-means clustering

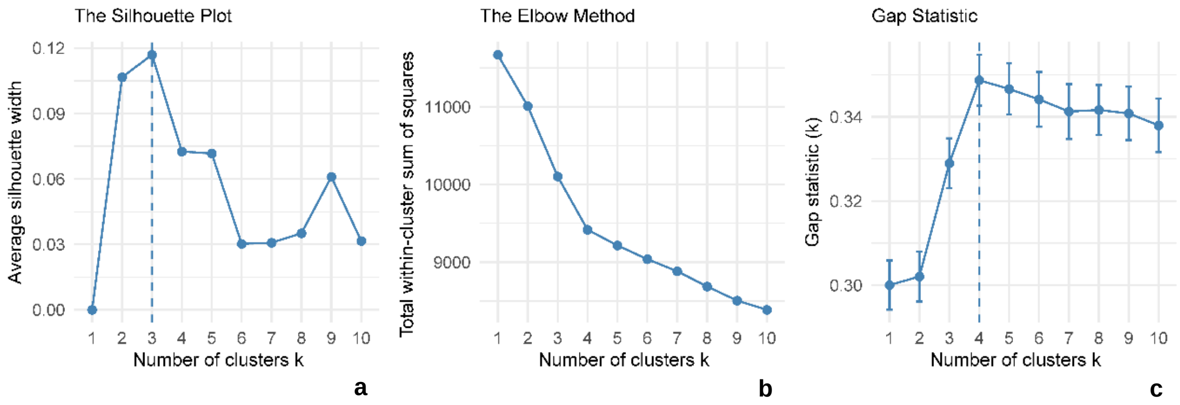


**Supplementary Fig. 8 Graphical output of methods used for determining the optimal number of clusters in the panel from a principal component analysis (PCA) using 622 dominant markers. (a)** Silhouette plot. **(b)** Elbow method. **(c)** Gap statistic

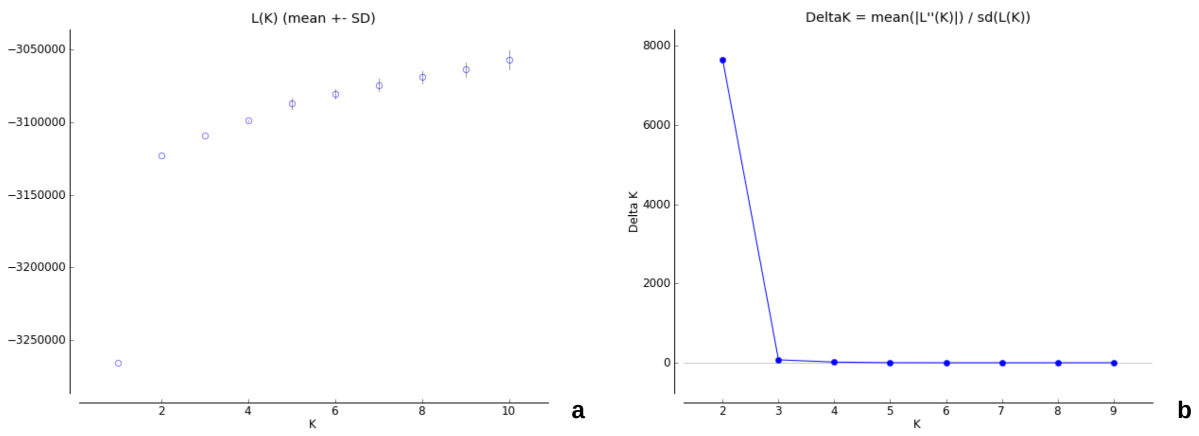


**Supplementary Fig. 9 Graphical output of methods used for the determination of the optimal number of genetic clusters in the panel based on the STRUCTURE Bayesian clustering analysis using 622 dominant markers. (a)** Mean LnP(D) for K = 1-10. **(b)** Mean ΔK for K = 1-10

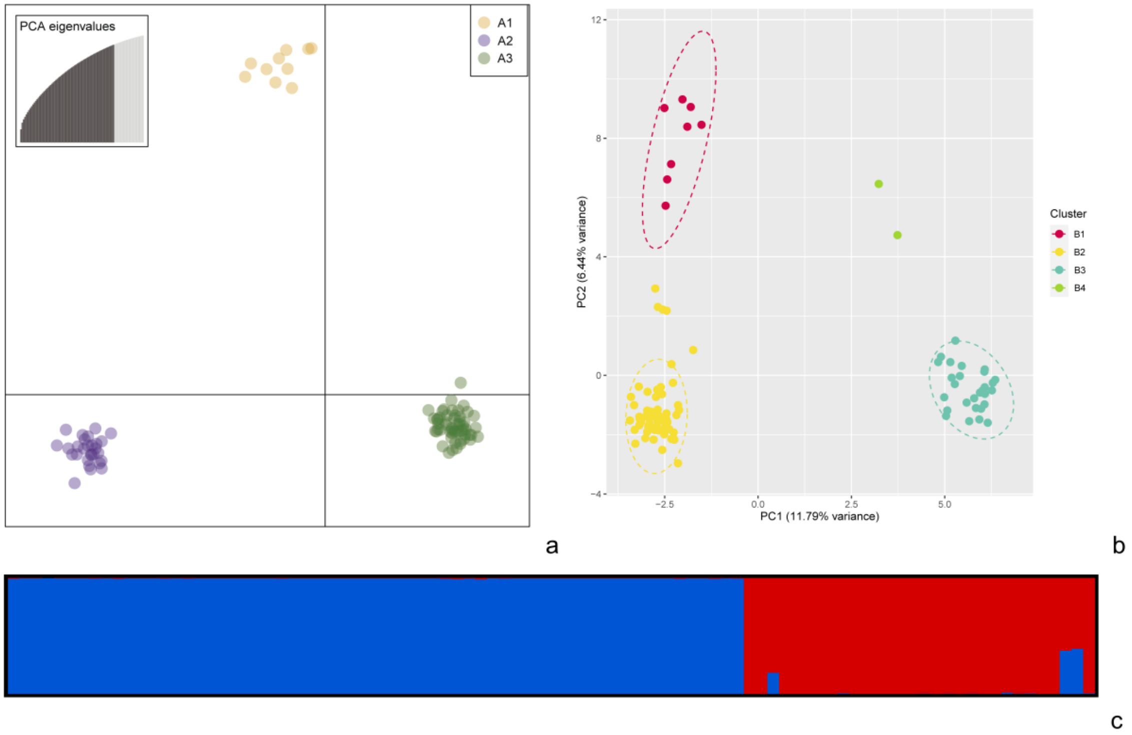


**Supplementary Fig. 10 Graphical outputs of the three clustering analyses employed to evaluate the genetic structure for 93 genotypes of the panel using 662 dominant markers. (a)** Plotting of the panel onto the first two linear discriminants from a discriminant analysis of principal components (DAPC). A division into three clusters (A1, A2 and A3) was considered. **(b)** Projection of the panel onto the first two principal components (PCs) from a principal component analysis (PCA). Genotypes are divided into four clusters (B1, B2, B3 and B4) as indicated by a k-means clustering analysis. Ellipses represent a 95% confidence interval. **(c)** Bayesian clustering performed on STRUCTURE. Each genotype is represented by a vertical bar split in accordance with the admixture coefficients for two clusters (C1 in dark blue and C2 in red). Genotypes are ordered according to ID number (see Supplementary Table 1)

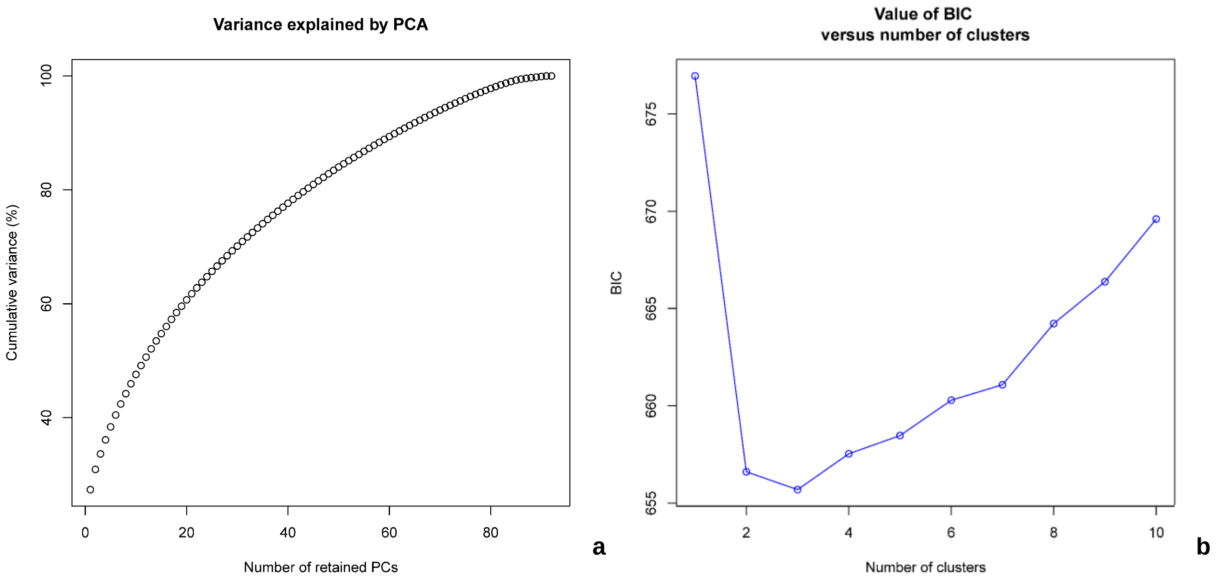


**Supplementary Fig. 11 Graphical output of methods employed for the determination of the optimal number of genetic clusters in the panel based on a discriminant analysis of principal components (DAPC) using 70,888 codominant markers. (a)** Cumulative variance explained by the principal components in the analysis. **(b)** Bayesian information criterion (BIC) versus number of clusters in k-means clustering

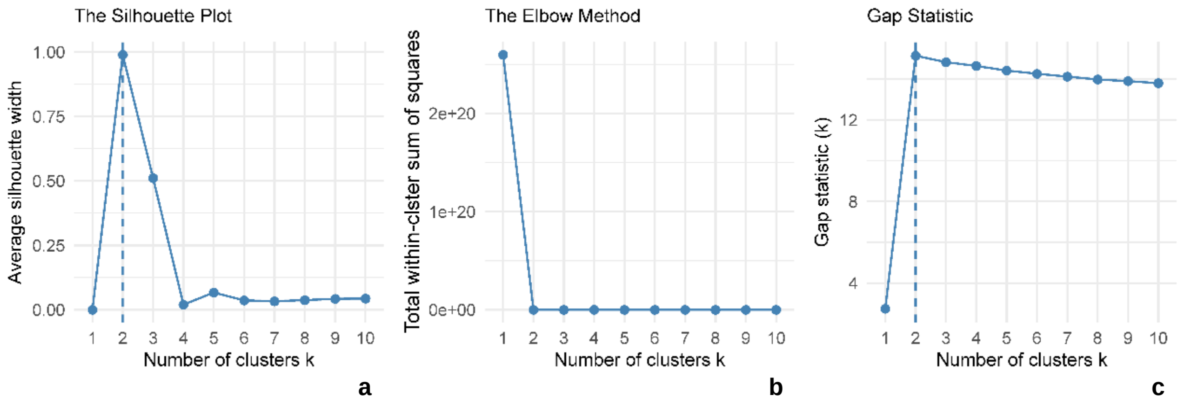


**Supplementary Fig. 12 Graphical output of methods used for determining the optimal number of clusters in the panel from a principal component analysis (PCA) using 70,888 codominant markers. (a)** Silhouette plot. **(b)** Elbow method. **(c)** Gap statistic

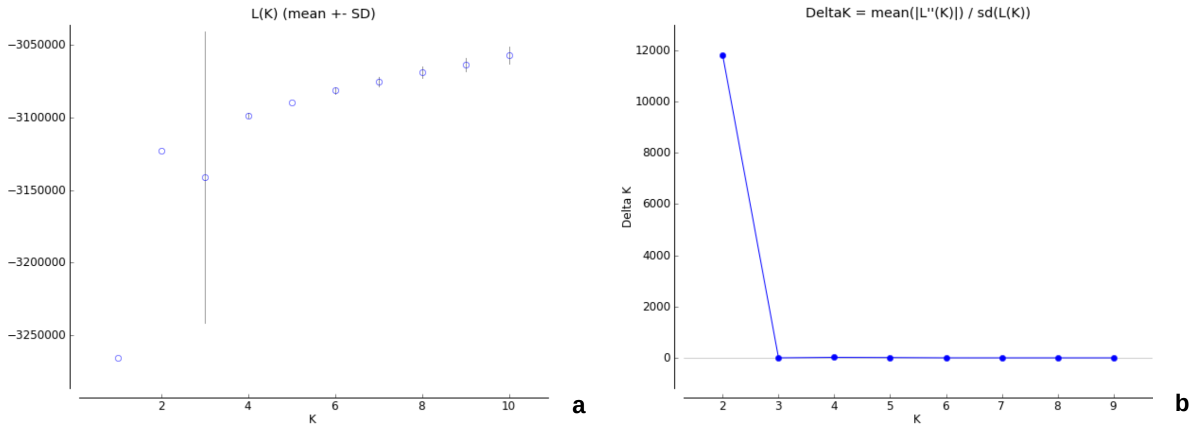


**Supplementary Fig. 13 Graphical output of methods used for the determination of the optimal number of genetic clusters in the panel based on the STRUCTURE Bayesian clustering analysis using 7,000 codominant markers. (a)** Mean LnP(D) for K = 1-10. **(b)** Mean ΔK for K = 1-10

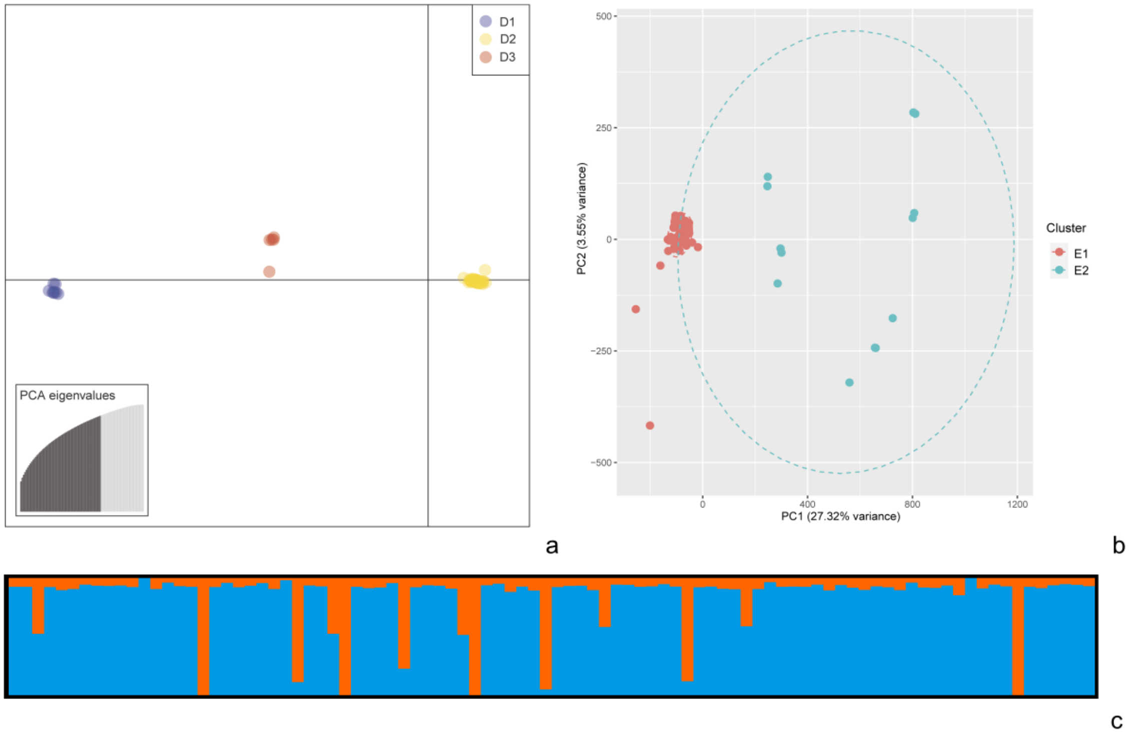


**Supplementary Fig. 14 Graphical outputs of the three clustering analyses employed to evaluate the genetic structures for 92 genotypes in the panel using codominant markers. (a)** Plotting of the panel onto the first two linear discriminants from a discriminant analysis of principal components (DAPC) using 70,888 single-nucleotide polymorphisms (SNPs) and insertions and deletions (indels). A division into three clusters (D1, D2 and D3) was considered. **(b)** Projection of the panel onto the first two principal components (PCs) from a principal component analysis (PCA) using 70,888 SNPs and indels. The genotypes were divided into two clusters (E1 and E2) as indicated by k-means clustering analysis. Ellipses represent a 95% confidence interval. **(c)** Bayesian clustering performed on STRUCTURE using 7,000 SNP and indel markers. Each genotype is represented by a vertical bar that is split in accordance with the admixture coefficients for two clusters (F1 in blue and F2 in orange). The genotypes are ordered according to ID number (see Supplementary Table 1)

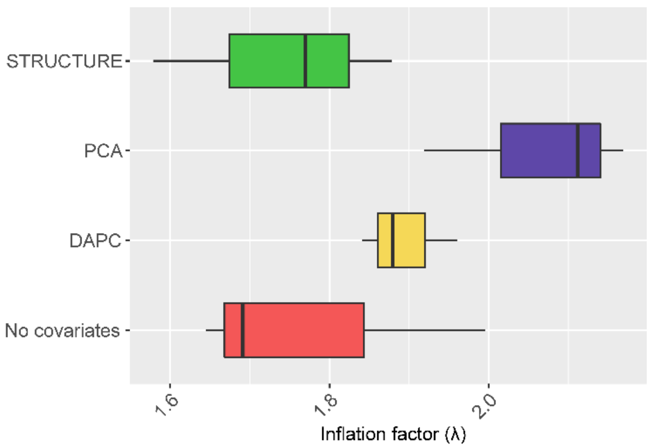


**Supplementary Fig. 15 Genomic inflation factor (λ) of the fixed and random model circulating probability unification (FarmCPU) models for the three SCYLV resistance traits analyzed, including different population structure matrices as covariates**. Abbreviations: DAPC, discriminant analysis of principal components; PCA, principal components analysis

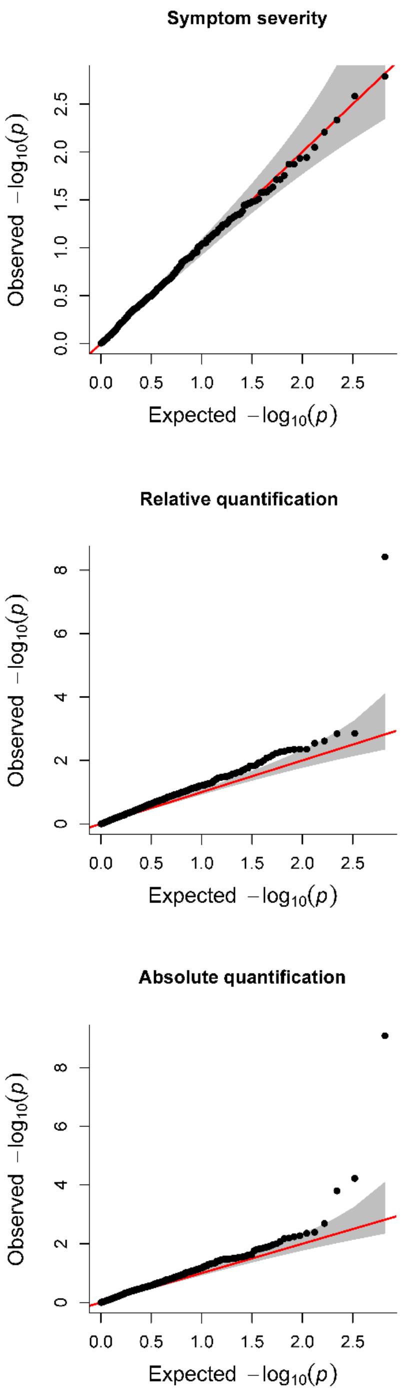


**Supplementary Fig. 16 Quantile-quantile (Q-Q) plots generated in the fixed and random model circulating probability unification (FarmCPU) analyses using the best linear unbiased predictor (BLUP) values of each of the three analyzed traits and 622 dominant markers**

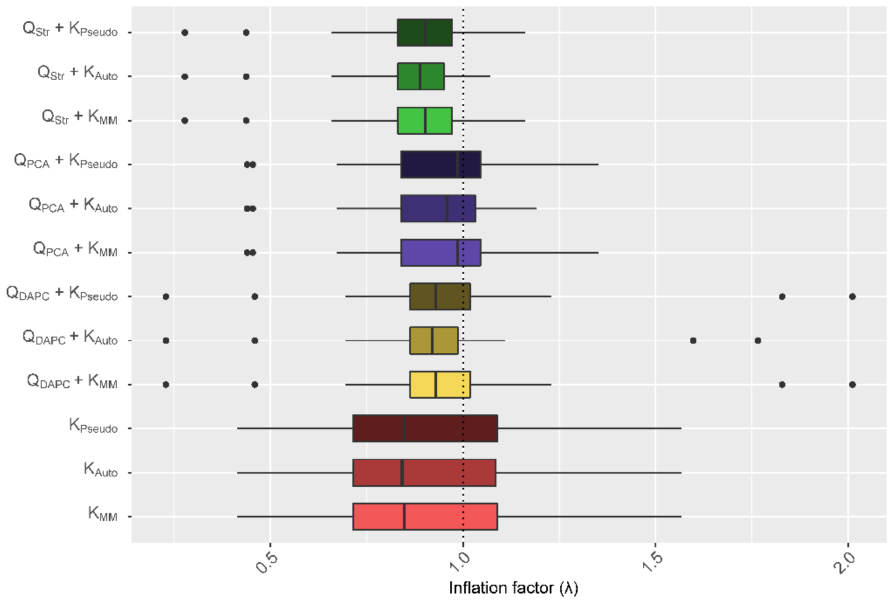


**Supplementary Fig. 17 Genomic inflation factor (λ) of twelve association models for the three SCYLV resistance traits analyzed, including different population structure (Q) and kinship (K) matrices as covariates.** Abbreviations: Auto, complete autopolyploid model; DAPC, discriminant analysis of principal components; MM, $MM^T$ realized relationship model; PCA, principal components analysis; Pseudo, pseudodiploid model; Str, STRUCTURE Bayesian clustering

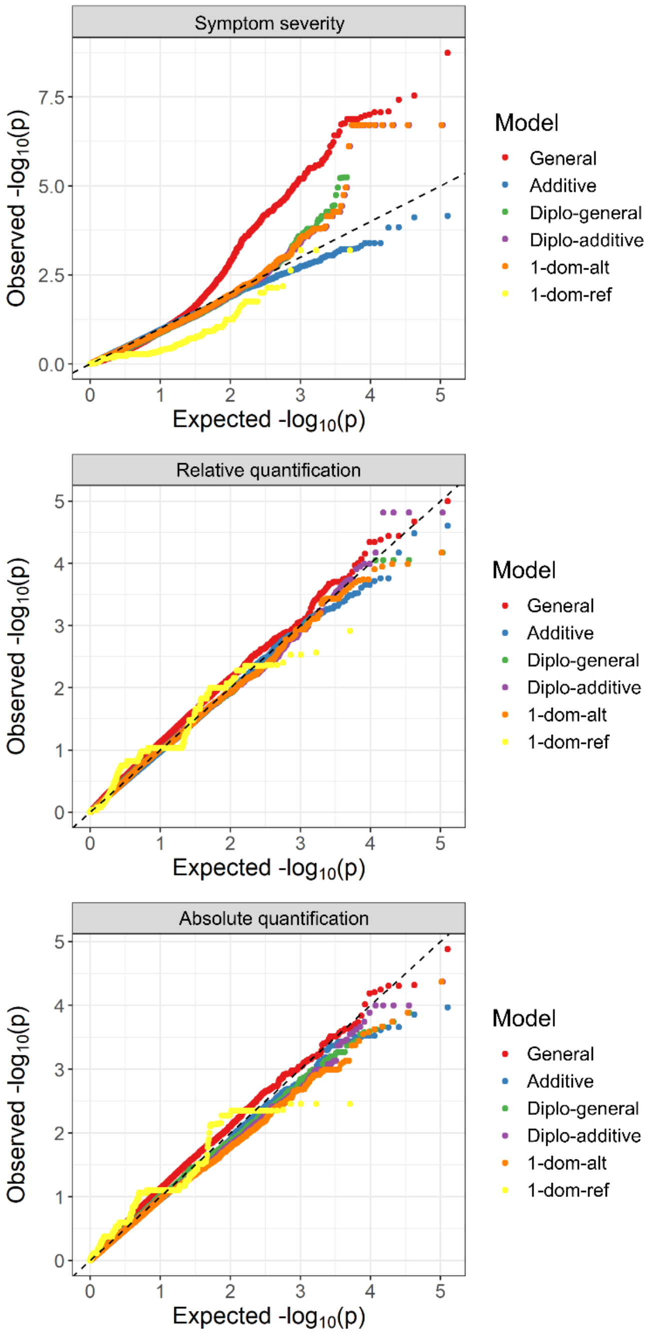


**Supplementary Fig. 18 Q-Q plots generated in the association analysis using the best linear unbiased predictor (BLUP) values of the three SCYLV resistance traits analyzed.** A realized relationship kinship matrix and the first three PCs from a PCA were used as covariates. Six different models were tested: general, additive, simplex dominant reference (1-dom-ref), simplex dominant alternative (1-dom-alt), diploidized general (diplo-general) and diploidized additive (diplo-additive)

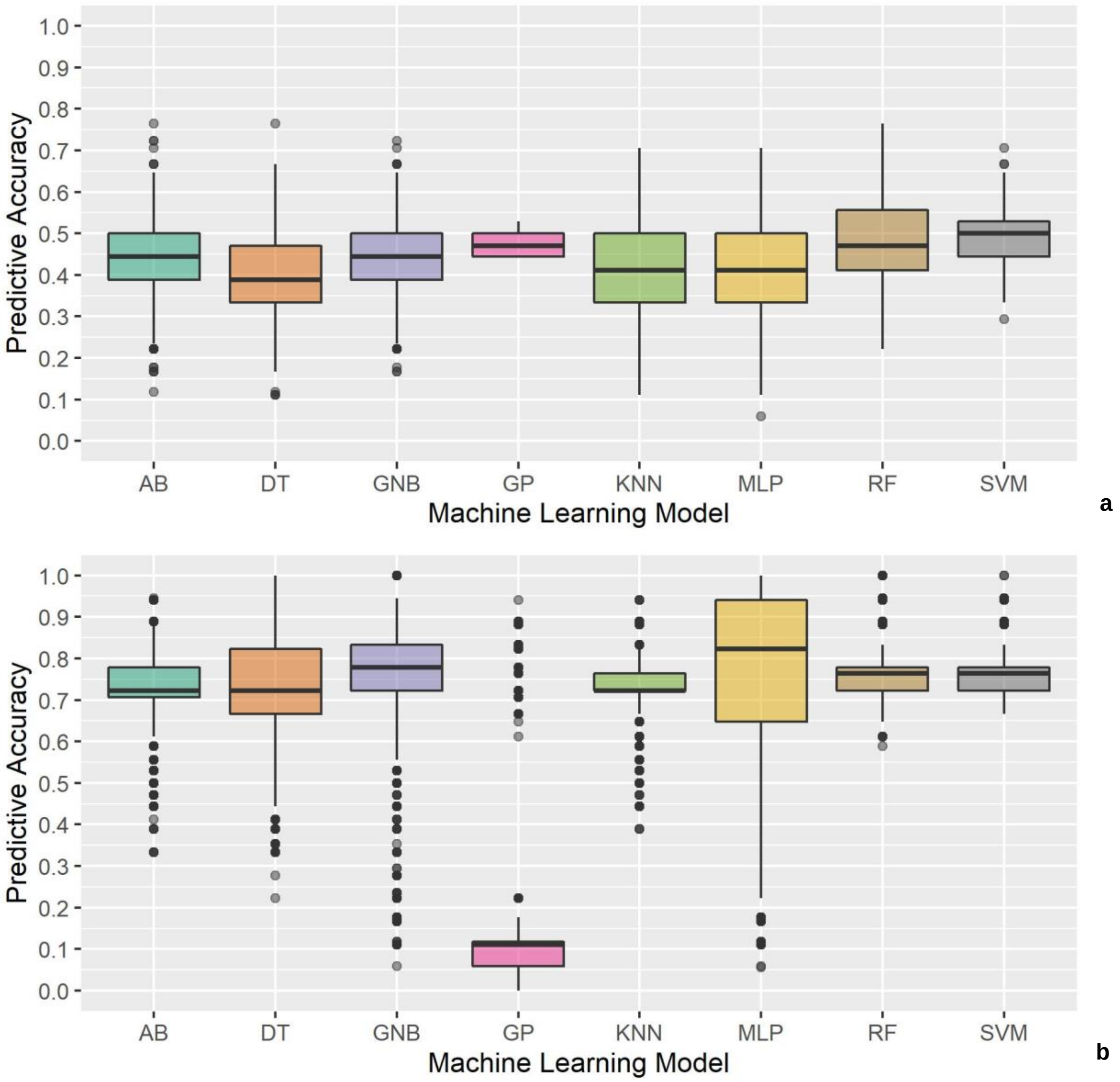


**Supplementary Fig. 19 Distribution of predictive accuracies of each machine learning approach employed to predict clusters associated with SCYLV resistance. (a)** Prediction of clustering by SCYLV titer determined by RT-qPCR (Q). **(b)** Prediction of clustering by SCYLV titer and SCYL symptom severity (SQ). The machine learning models tested were adaptive boosting (AB), decision tree (DT), Gaussian naive Bayes (GNB), Gaussian process (GP), K-nearest neighbor (KNN), multilayer perceptron neural network (MLP), random forest (RF) and support vector machine (SVM)

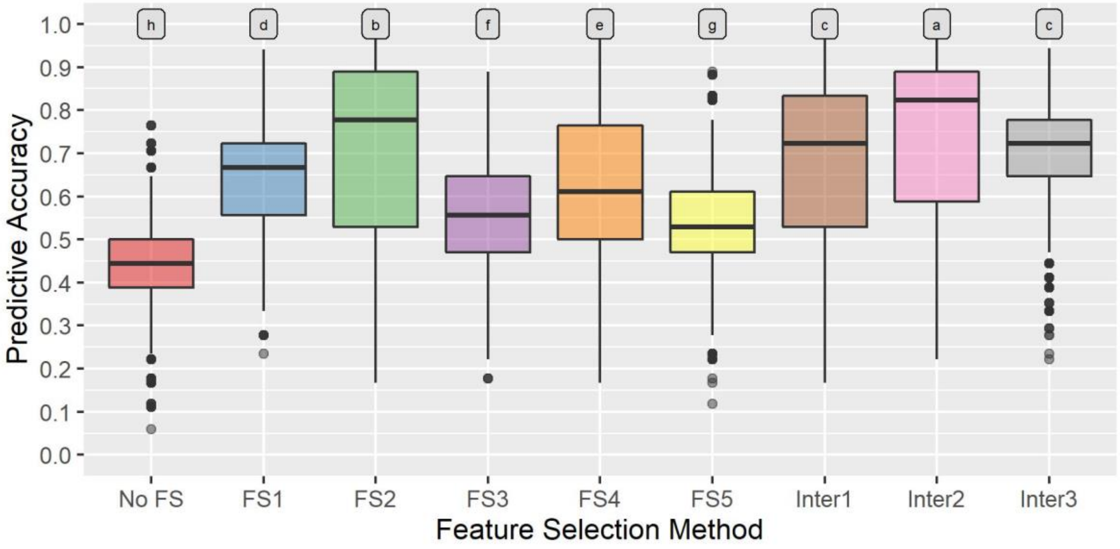

a

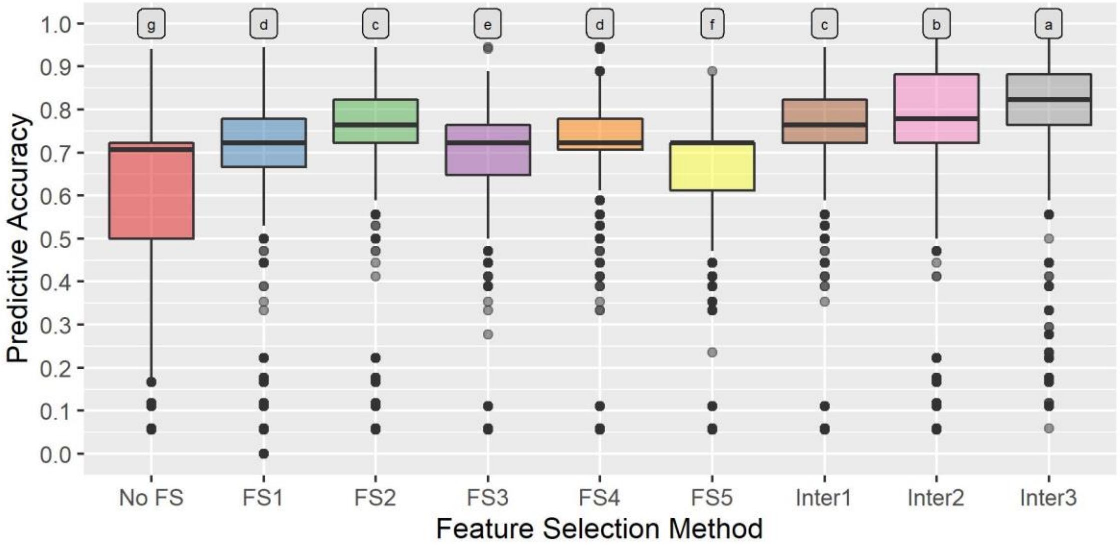

b

**Supplementary Fig. 20 Distribution of predictive accuracies of all machine learning approaches employed to predict clusters associated with SCYLV resistance using each marker dataset obtained by feature selection (FS) strategies. (a)** Prediction of clustering by SCYLV titer determined by RT-qPCR (Q). **(b)** Prediction of clustering by SCYLV titer and SCYL symptom severity (SQ). FS methods tested were gradient tree boosting (FS1), L1-based FS through a linear support vector classification system (FS2), extremely randomized trees (FS3), univariate FS using ANOVA (FS4) and RF (FS5), in addition to the intersection of markers selected by at least two of all FS methods (Inter1), the intersection of at least three of the three best FS methods (Inter2) and the intersection of the three best FS methods (Inter3). Letters above boxplots indicate significantly different groups identified by ANOVA followed by Tukey's test
